# Disruption of *grin2A*, an epilepsy-associated gene, produces altered spontaneous swim behavior in zebrafish

**DOI:** 10.1101/2025.05.09.651809

**Authors:** Vera Abramova, Eni Tomovic, Bohdan Kysilov, Miloslav Korinek, Mark Dobrovolski, Barbora Hrcka Krausova, Klevinda Fili, Fatma Elzahraa S. Abdel Rahman, Paulina Bozikova, Jiri Cerny, Tereza Smejkalova, Ales Balik, Ladislav Vyklicky

**Author notes:** **Corresponding authors**: Prof. Ladislav Vyklicky, MD, PhD Institute of Physiology, Czech Academy of Sciences Videnska 1083, 142 00 Prague 4, Czech Republic Ales Balik, PhD Institute of Physiology, Czech Academy of Sciences BIOCEV, Prumyslova 595, 252 50 Vestec, Czech Republic.

## Abstract

N-methyl-D-aspartate receptors (NMDARs) control synaptic plasticity and brain development in a manner determined by receptor subunit composition. Pathogenic variants in *GRIN2A* gene, encoding the NMDAR GluN2A subunit, can cause gain or loss of function of receptors containing the affected subunit, and are associated with intellectual disability and epilepsy in patients. While *in-vitro* studies of recombinant receptors have yielded some insights, animal experimental models are essential to better understand the relationship between the molecular pathology of the variants and the disease. Here we introduce a zebrafish model of GluN2A loss of function to study system-level effects of zebrafish *grin2Aa* and *grin2Ab* gene deletion. Our electrophysiological analysis revealed functional differences between receptors containing zebrafish GluN2Aa/b and GluN2Bb paralogs comparable to mammalian receptors containing GluN2A *vs.* GluN2B subunits. Both *grin2Aa^−/−^* and *grin2Ab^−/−^*, as well as double-knockout *grin2A^−/−^* zebrafish larvae showed increased locomotor activity in a novel environment. Proteomic analysis suggested that the relative proportion of GluN2B-containing NMDARs may be increased in *grin2A* mutant fish. Our results highlight fundamental similarities between zebrafish and mammalian NMDAR signaling and validate the use of zebrafish as a model organism to study the neurodevelopmental role of NMDARs. The newly created transgenic zebrafish strains complement the rodent models of GluN2A loss of function and can be used for high-throughput testing of pharmacological or genetic treatment strategies for patients with *GRIN2A* gene variants.

## INTRODUCTION

Brain function depends on synaptic transmission, ∼80% of which is mediated by presynaptically released glutamate that activates postsynaptic ionotropic receptors, including N-methyl-D-aspartate receptors (NMDAR) (Hansen et al., 2021). NMDAR-mediated signaling is critical for synaptic plasticity (Dingledine et al., 1999; Lynch, 2004; Traynelis et al., 2010; Huganir and Nicoll, 2013), brain development and learning and memory (Hansen et al., 2021). NMDARs are heterotetrameric complexes composed of two GluN1 and two GluN2(A-D)/GluN3(A­B) subunits (Traynelis et al., 2010). Genome-wide association studies have linked *de-novo* and inherited missense and nonsense/frameshift variants in *GRIN* genes, which encode NMDAR subunits, to a range of neurodevelopmental disorders—including intellectual disability (ID), developmental delay, autism spectrum disorder (ASD), and epilepsy (Soto et al., 2014; Burnashev and Szepetowski, 2015; Hu et al., 2016; XiangWei et al., 2018; Garcia-Recio et al., 2021; Benke et al., 2021). While many variants have yet to be functionally characterized, common molecular effects include altered surface expression and loss- or gain-of-function changes (Yuan et al., 2014; Swanger et al., 2016; Chen et al., 2017; Ogden et al., 2017; Vyklicky et al., 2018; Amin et al., 2021; Korinek et al., 2024). The relationship between the molecular pathology of the variants and the disease is poorly understood. In the case of disease-associated variants of *GRIN2A*, loss of function of GluN2A-containing NMDARs or reduced availability of GluN2A protein may paradoxically lead to circuit hyperexcitability resulting in epilepsy (Swanger et al., 2016; Camp et al., 2023).

In recent years, zebrafish (*Danio rerio*) have emerged as a powerful alternative to mammalian models for studying brain disorders and for high-throughput drug screening (Bruni et al., 2016; Leung and Mourrain, 2016; Zoodsma et al., 2022; Zoodsma et al., 2020; Abramova et al., 2023). The zebrafish model has several advantages: low maintenance cost, high fecundity, rapid development, and amenability to genetic manipulation. Although zebrafish are simple organisms, they share high genetic homology with humans (Howe et al., 2013; Barbazuk et al., 2000).

Basic neurophysiology is conserved in vertebrates, and zebrafish neurotransmitters and receptors, including the NMDAR subtype, are similar to those found in mammals (Panula et al., 2010; Cox et al., 2005). Due to teleost-specific genome duplication, zebrafish possess two paralogs for many genes (Meyer and Schartl, 1999; Furutani-Seiki and Wittbrodt, 2004), including 13 putative genes encoding NMDAR subunits compared to 7 genes in humans (Horzmann and Freeman, 2016). Functional studies have shown that deletion of *GRIN1* or *GRIN2B* paralogs (e.g., *grin1a*, *grin1b*, *grin2Ba*, *grin2Bb*), alters behavior (Zoodsma et al., 2020; Zoodsma et al., 2022).

In the present study, we examined the molecular and behavioral consequences of deleting *grin2Aa* and *grin2Ab*, zebrafish paralogs of *GRIN2A*. We show that: (1) zebrafish NMDARs exhibit subunit-dependent deactivation kinetics, with receptors containing the zGluN2A subunit deactivating more rapidly than those containing the zGluN2B subunits; (2) *grin2Aa* and *grin2Ab* mRNA is developmentally regulated; (3) knockout of *grin2Aa* or *grin2Ab* or both increases locomotor activity, unlike *grin2Bb* knockout, which shows no change; and (4) GluN2B-containing NMDARs are relatively enriched in *grin2A-*mutant fish, while the total NMDAR abundance remains unchanged. Together, these findings support the utility of zebrafish as a model for *GRIN*-related neurodevelopmental disorders and complement rodent-based studies on GluN2A loss-of-function.

## METHODS

### Transfection and maintenance of HEK cells

Human embryonic kidney 293T (HEK293T) cells (American Type Culture Collection, ATCC No. CRL1573, Rockville, MD, USA) were cultured in Opti-MEM I (Invitrogen, Carlsbad, CA, USA) with 5% fetal bovine serum (PAN Biotech, Aidenbach, Germany) at 37°C in 5% CO_2_. One day before transfection, cells were plated in 24-well plates at a density of 2×10^5^ cells per well. The next day, cells were transfected with pcDNA3.1+ expression vectors containing cDNA encoding the zebrafish wild-type NMDA receptor subunit zGluN1a (ENSDART00000102368.5), zGluN1b (ENSDART00000034849.9), zGluN2Aa (ENSDART00000142141.5), zGluN2Ab (ENSDART00000103652.5), zGluN2Ba (XM_009299013.2), and zGluN2Bb (ENSDART00000143664.3) (purchased from GenScript, The Netherlands), and green fluorescent protein (GFP; pQBI 25, Takara, Tokyo, Japan). Briefly, equal amounts (0.2 µg) of cDNAs encoding zGluN1, one of paralog zGluN2A or zGluN2B and GFP were mixed with 0.6 μL MATra-A Reagent (IBA, Göttingen, Germany) and added to confluent HEK293T cells cultured in 24-well plates. After trypsinization, the cells were resuspended in Opti-MEM I containing 1% fetal bovine serum supplemented with 20 mM MgCl_2_, 3 mM kynurenic acid, 1 mM D,L-2-amino-5-phosphonovaleric acid, and 2 μM ketamine and plated on 30 mm collagen and poly-L-lysine-coated glass coverslips. Transfected cells were visualized by GFP epifluorescence. Amino acids are numbered according to the full-length protein, including the signal peptide with the initiating methionine as number 1.

### Whole-cell patch-clamp electrophysiology

Experiments were performed 24-36 hours after the end of transfection in cultured HEK293T cells co-transfected with genes encoding zGluN1, zGluN2, and GFP. Whole-cell voltage-clamp current recordings were performed at room temperature (21-25°C) at a holding potential of ­60 mV using a patch-clamp amplifier (Axopatch 200B; Molecular Devices, Sunnyvale, CA, USA) after 80-90% compensation for capacitance and series resistance (<10 MΩ). Data were collected (sampled at 10 kHz and low-pass filtered at 2 kHz) and analyzed using pClamp 10 (Molecular Devices). Patch pipettes (3­5 MΩ) were filled with intracellular solution containing (in mM): 120 gluconic acid, 15 CsCl, 10 BAPTA, 10 HEPES, 3 MgCl_2_, 1 CaCl_2_, and 2 ATP­Mg salt (pH adjusted to 7.2 with CsOH). Extracellular solution (ECS) contained the following (in mM): 160 NaCl, 2.5 KCl, 10 HEPES, 10 glucose, 0.2 ethylenediaminetetraacetic acid, and 0.7 CaCl_2_ (pH adjusted to 7.3 with NaOH). Special precautions were taken to reduce the background concentration of glycine in experiments examining the glycine dose-response relationship, which involved using ultrapure chemicals and HPLC water (Merck). Agonist applications were performed using a microprocessor-controlled multi-barrel fast perfusion system with a solution exchange rate of τ ∼12.0 ms (Vyklicky et al., 2018).

### Zebrafish maintenance and housing

Zebrafish (*Danio rerio*) were handled and maintained in accordance with Directive 2010/63/EU on the protection of animals used for scientific purposes, and national, and institutional guidelines. Fish were maintained according to published protocols (Westerfield, 2000) and zebrafish husbandry recommendations (Alestrom et al., 2020), and housed in a ZebTEC aquatic system (Tecniplast) at 28°C in the CZ-OPENSCREEN fish facility (Institute of Molecular Genetics CAS, Prague). The fish were fed daily with live brine shrimp and dried flake food (Skretting, Tooele, USA). They were maintained on a 14 h light:10 h dark cycle (lights on at 8:00; lights off at 22:00) (see Abramova et al., 2023 for details). Embryos and larvae were raised in E3 medium (in mM): 5 NaCl, 0.17 KCl, 0.33 CaCl_2_, 0.33 MgSO_4_ with 0.0001% methylene blue at 28°C and under the same light conditions as adult fish.

The mutant *grin2Aa* zebrafish (ID: ZDB-ALT-130411-3272, allele ref. sa14573) were purchased from Zebrafish International Resource Center (Eugene, OR, USA) (Dooley et al., 2013; Kettleborough et al., 2013). They were outcrossed to AB wild-type fish. The zebrafish *grin2Ab* and *grin2Bb* mutant strains were generated using CRISPR/Cas9 gene targeting of fertilized embryos on the AB background.

### CRISPR-Cas9 gene targeting and embryo microinjections

sgRNA (TGGAGGAAGCCCGTTCGCTA, purchased from Sigma) targeting exon 4 of *grin2Ab* was designed using CRISPRscan (Moreno-Mateos et al., 2015). 5 µl of injection mix included: 0.3 µl sgRNA (1000 ng/µl), 0.4 µl Cas9-EGFP protein (1.7 mg/ml), 0.4 µl 1x dilution buffer (from CAS9GFPPRO kit, Sigma), 2 M KCl, 0.5 µl phenol red (Sigma), 0.5 µl 10X NEB3.1 buffer (New England Biolabs, USA), and 2.4 µl RNAse-free H_2_O. Single-cell stage fertilized wild type embryos (AB background) were injected with 1 nl of injection mix containing 60 pg of sgRNA and 400 pg of Cas9-EGFP protein. Injected embryos were incubated at 34°C for 4 h and then transferred to fresh medium at 28°C and dechorionated. The next day, 10-20 embryos from each injection batch (200 eggs) were tested for the presence of premature stop codon by PCR using primers flanking the sgRNA target site (see Table S1 for primers used). The PCR products obtained were separated on native 5% PAGE. The presence of a premature stop codon appeared as extra bands. The positive batch was reared until adulthood. Adult fish were tested for the presence of a premature stop codon (mosaicism) by fin clip genotyping and, if positive, outcrossed to the wild type fish (AB strain) to test for germline transmission. Out of 48 tested adult fish, one transmitted a 16 bp deletion resulting in a frame shift and subsequent appearance of a premature stop codon in *grin2Ab*. Founder fish were outcrossed twice to the wild type fish (AB strain) to rule out non-target effects of CRISPR/Cas9 gene editing. The *grin2Bb* mutant strain was generated by the same approach using the following sgRNA AGGTATAACCGTAACCAGTG targeting exon 4 of *grin2Bb*, producing a 1 bp deletion resulting in a frameshift and a premature stop codon in *grin2Bb*.

### Zebrafish genotyping

After each behavioral experiment, the larvae were collected, and the genotype was determined. Genotyping was performed by direct sequencing (Eurofins Genomics, Germany) of PCR amplicons using primers shown in Table S1. For more details on genotyping, sequencing, and primers see Table S1.

### Whole-mount RNA in-situ hybridization (ISH)

ISH was performed as described (Vauti et al., 2020). Briefly, templates for *grin2Aa* and *grin2Ab* RNA probe synthesis were amplified from zebrafish cDNA (primers shown in Table S1) and cloned into pGEM-T Easy vector (Promega, USA). Positive colonies were selected by IPTG/X-gal (blue-white) screening and the presence of the insert was verified by sequencing (Eurofins Genomics, Germany). Plasmids containing the insert with an antisense orientation under the regulation of the T7 promoter were selected for the synthesis of antisense probes. All plasmids for *in-vitro* sense and antisense probe synthesis were isolated using the NucleoBond Xtra Midi or Maxi kit (Macherey Nagel, Germany) from bacterial cultures grown overnight.

For probe *in-vitro* synthesis, plasmids were linearized, separated and excised from 1% agarose gel, and purified using the Monarch DNA Gel Extraction Kit (New England Biolabs). DIG RNA Labeling Mix Kit (Roche, Switzerland) was used for digoxigenin-UTP incorporation into the RNA probe. The RNA probe was purified by 8 M LiCl precipitation. Quality control of synthesized RNA probes was performed in denaturing (formaldehyde) 1% agarose gel. Wild type (AB strain) larvae were collected at day 1, 3, or 5 post fertilization (dpf), depigmented in 0.003% phenylthiol urea, fixed in 4% paraformaldehyde, dehydrated through the series of ethanol, and stored at -20°C in methanol. On the day of the experiment, larvae were rehydrated through the series of ethanol, permeabilized in 80% acetone, bleached, and hybridized with the desired RNA probes at 65°C in a water bath for 60 hours. After RNA hybridization, larvae were washed and incubated with anti-digoxigenin-alkaline phosphatase Fab fragments (1:2000) (Roche, Switzerland). The larvae were then washed in PBS (2 days) and finally hybridization signal was developed with BM Purple (Roche). The images of larvae were acquired on the Zeiss Stereo Discovery V20 microscope and then processed using CorelDRAW Graphics Suite.

### RNA isolation and RT-qPCR expression analysis

Total RNA was extracted from the individual heads of zebrafish wild-type and mutant larvae (6 dpf) and from the optic tectum and ventral structures, including thalamic and hypothalamic nuclei, of adult fishes (three months old). Heads or brain tissue were frozen on dry ice immediately after dissection and stored at -80°C until RNA isolation. The tissue samples were homogenized in TRI Reagent (Merck), and total RNA was extracted according to the manufacturer’s protocol. The RNA pellet was resuspended in 20 µl of RNase-free water and the concentration of RNA was measured using a Nano Drop instrument (Thermo Scientific). We used 500 ng of total RNA for cDNA synthesis. 250 ng of random primers (Promega) was combined with total RNA, incubated at 70°C for 5 min and then cooled on ice for 2 min. Subsequently, cDNA was synthetized (at 37°C, for 90 min) in a 25 µl reaction containing 200 U of M-MLV Reverse Transcriptase (Promega). The obtained cDNA was additionally diluted by 25 µl of PCR grade water and stored at 4°C until RT-qPCR analysis.

For RT-qPCR, the cDNA samples (1.5 μl) were added into RT-qPCR reaction mixture (final volume 20 μl) containing RT-qPCR 2x Sybr Master Mix (P553, Top-Bio, Czech Republic) and specific primers as described previously or newly designed using an on-line tool (Eurofins Genomics) (Table S1). All RT-qPCR reactions were performed in triplicates and run on a LightCycler 480 instrument (Roche Diagnostics). The mean of the crossing point (Ct values) obtained from RT-qPCR was normalized to the average level of two housekeeping genes *β-actin* and *elfa* (*elongation factor 1α*) and used for the analysis of relative gene expression (RQ values) by the ΔΔC_T_ method (Livak and Schmittgen, 2001; Taylor et al., 2019).

### Mass spectrometry analysis

The optic tectum and ventral structures, which include thalamic and hypothalamic nuclei, of adult zebrafish from wild type (AB strain) controls, *grin2Aa^−/−^* (four biological replicates), and *grin2Ab^−/−^* fish (four biological replicates) were lysed by multiple rounds of sonication in 100 mM triethylammonium bicarbonate, pH 7.0, containing 2% sodium deoxycholate. The samples were then processed as previously described (Candelas Serra et al., 2024).

Peptides were separated and analyzed on an UltiMate 3000 RSLCnano system coupled to an Orbitrap Fusion Tribrid mass spectrometer (both Thermo Scientific). An EASY-Spray column (75 µm x 50 cm) (Thermo Scientific) was used for peptide separation in combination with Acclaim PepMap300 trap column (Thermo Scientific). 180-minute linear gradient from 5% acetonitrile to 35% acetonitrile was used for separation at a flow rate of 300 nl/min. (McAlister et al., 2014).

Raw data were processed in ProteomeDiscoverer 2.5 (Thermo Scientific). Tandem mass tag reporter ion ratios were used to estimate of the relative amount of each protein. The search was performed against *Danio rerio* and the common contaminant database. The modification was set: Tandem mass tag-pro label at peptide N-terminus and lysine (unimod nr:2016), cysteine caramidomethylation (unimod nr:39) as static; methionoine oxidation (unimod nr:1384), and protein N-terminus acetylation (unimod nr:1) as variable. Proteins and peptides were filtered to 1% false discovery rate (FDR).

Data were normalized to total peptide abundance, and all subsequent analyses were performed in Perseus software (Tyanova and Cox, 2018). We filtered out contaminants and logarithmized intensities (binary logarithms) and filtered out proteins with insufficient number of valid quantification values (only those with at least 3 values in at least one group were retained). Student’s t-test with permutation-based FDR correction was used to evaluate significantly altered proteins at the 5% FDR level.

The “proteomic ruler” approach was applied to estimate the copy abundance number (CAN) of individual proteins per cell, as described by (Wisniewski et al., 2014). This method is based on the mass spectrometry signal of histones, which is proportional to the DNA content in the sample and, consequently, to the number of cells. The Proteomics Ruler plugin in Perseus was utilized for CAN estimation, with scaling set to the total protein amount. The relative CAN was then calculated using the CAN of GluN1a in *grin2A^+/+^*fish as a reference.

### Calculation of proportions of NMDA receptors with various subunit compositions

Mass spectrometry provided copy abundance numbers (C) for the subunits of NMDA receptors GluN1a, GluN2Aa, GluN2Ab, GluN2Ba and GluN2Bb (*C*_1_, *C*_2Aa_, *C*_2Ab_, *C*_2Ba_, *C*_2Bb_). To calculate the proportion of NMDA receptors with a particular subunit composition, we assumed that every NMDAR tetramer consists of two obligatory GluN1 subunits and the two places for GluN2 subunits are randomly occupied by the above mentioned GluN2 subunit paralogues.

Therefore, the relative number of tetramers with 1a-1a-2Aa-2Aa subunit composition is proportional to (*C*_2Aa_)^2^. The relative number of tetramers with 1a-1a-2Aa-2Ab subunit composition is proportional to 2*C*_2Aa_*C*_2Ab_(this expression contains the factor of 2 since combinations of 2Aa-2Ab and 2Ab-2Aa provide receptors with the same subunit composition). This is how we calculated relative numbers of tetramers of all GluN2 subunit compositions (altogether GluN1 together with GluN2Aa, GluN2Ab, GluN2Ba and GluN2Bb can assemble to 10 subunit compositions). Out of all assembled receptors, the proportion of receptors with 1a-1a-2Aa-2Aa subunit composition (*P*_2Aa-2Aa_) is equal to

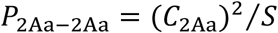

where *S* is the sum of relative numbers of all assembled receptors:

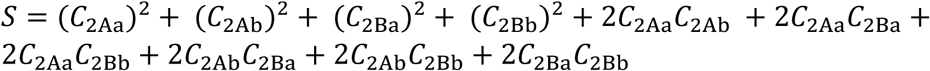

Similarly, out of all assembled receptors, the proportion of

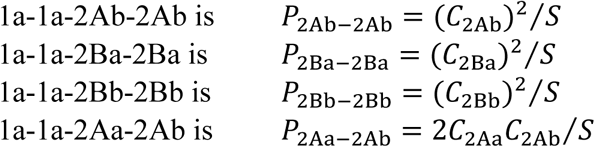

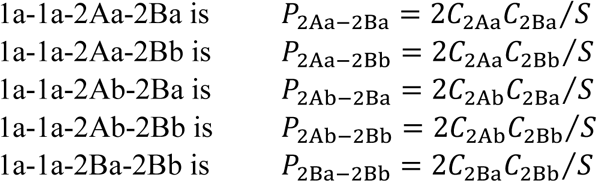

Our electrophysiology measurements showed that receptors containing one or two copies of GluN2Ba (4 subunit compositions) are not functional. Six remaining subunit compositions provide functional receptors. Therefore, out of all functional receptors, the proportion of receptors with 1a-1a-2Aa-2Aa subunit composition (*R*_2Aa-2Aa_) is equal to

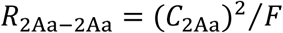

Where F is the sum of relative numbers of functional receptors:

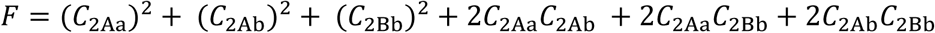

Similarly, out of all functional receptors, the proportion of

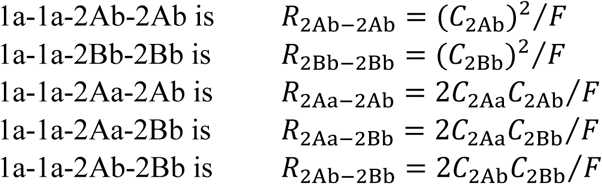

### Assay of the locomotor activity

All experiments were performed in groups of 20 zebrafish larvae at 6 days post-fertilization (dpf). Larvae were individually transferred from sibling-containing Petri dishes (maximum 50 larvae per 100 mm Petri dish) into custom-fabricated observation wells. Each well was a 10 mm x 10 mm x 3 mm (w x d x h) square compartment, separated by 2 mm thick septa made of poly-D,L-lactic acid and arranged in a 4 x 5 grid, 3D-printed using Original Prusa i3 MK3S printer (Prusa Research, Czech Republic). The grid was fixed to the bottom of a 100 x 20 mm polystyrene Petri dish (# 130182, Thermo Scientific). The dish was maintained at 28 ± 0.5°C under constant illumination (100 lux), and placed in a custom-made holder on an optically and vibration-isolated table. Locomotor activity was recorded for 1 hour immediately after larval transfer with experiments conducted between 14:00 and 19:00. Video monitoring and behavioral analysis were performed as previously described (Abramova et al., 2023). Briefly, recordings were captured at 50 frames per second and stored for post-hoc analysis. Each video frame was cropped into individual wells, and a ResNet-based deep learning model, trained on hand-annotated larval images, was used to identify larval position and orientation. Position was defined using two key points: one for the head between the eyes and one in the center of the swimming bladder. Larval movement tracking was performed using a custom Python script implemented with TensorFlow and Keras libraries.

Tracks were analyzed for the mean travel distance (MTD), mean bout distance (MBD), mean bout frequency (MBF), and thigmotaxis. The travel distance was defined as the distance a larva traveled over 10 min period, calculated as the sum of all movements recorded at 20 ms intervals (corresponding to the 50 fps video frame rate). To eliminate/remove system noise, a minimum movement threshold of 0.134 mm per 20 ms was applied. A bout was defined as a single movement or multiple movements separated by stillness longer than 140 ms. MBD was calculated as the average length of these movement bouts, and MBF as the total number of bouts within a 10 min interval. Thigmotaxis was assessed by measuring the relative time each larva spent in the central zone of the well during the same 10 min period. The central zone was defined as a virtual square measuring 5.5 x 5.5 mm centered within each 10 x 10 mm well. Behavioral assays were conducted on larvae from multiple independent crosses and different parental combinations to minimize the effects of genetic background.

### Experimental design and statistical analysis

Data were analyzed using the statistical software program Statgraphics Centurion 18 (Statgraphics Technologies, Inc., VA, USA). All reported averages are presented with their corresponding standard error of the mean (SEM). The number of replicates is indicated in the figure legends. If necessary, data were transformed using the power function to ensure a symmetric distribution and constant variance. The analysis of variance was performed using the general linear models function, with genotype as the between-subject factor and time bin as the within-subject factor for repeated measures ANOVA testing. The absolute value of the studentized residuals was calculated and compared to a threshold of 3, which was used to identify and exclude outliers. Throughout the study, *p* ≤ 0.05 was considered statistically significant. In cases where the *p*-value indicated a statistically significant difference, the subsequent analysis was conducted using the least significant difference (LSD) test or pairwise comparisons with the Bonferroni method (where multiple comparison correction was necessary) or the Dunnett’s method for multiple comparisons versus the control.

## RESULTS

### Zebrafish GluN2A paralogs: Homology and function

In mammals, NMDARs are tetrameric complexes composed of two GluN1 subunits (encoded by the *GRIN1* gene) and a combination of two GluN2 (GRIN2A–D) and/or GluN3 (GRIN3A– B) subunits. Each subunit includes a large extracellular region with an amino-terminal domain (ATD) and an agonist-binding domain (ABD), a transmembrane domain (TMD) comprising helices M1, M3, M4, and a reentrant M2 loop, and an intracellular C-terminal domain (CTD) (Hansen et al., 2021) (Fig. 1A). In zebrafish, all NMDAR subunit genes except *grin3A* exist as two paralogs, each sharing high sequence homology with its human ortholog (Cox et al., 2005).

**Figure 1.**
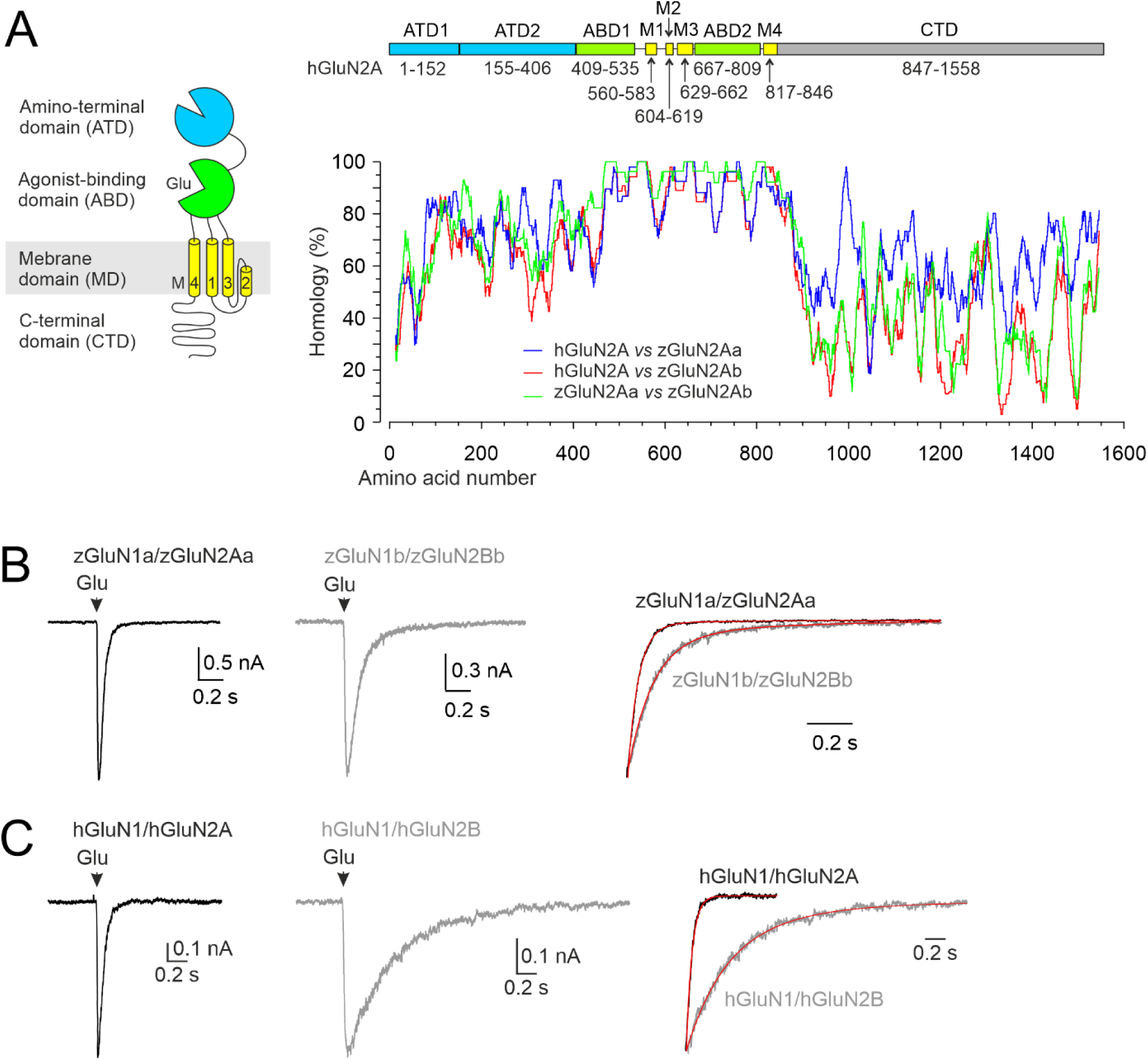
Protein sequence homology and functional properties of recombinant zebrafish NMDARs. A, Cartoon of the domains comprising the hGluN2A subunit: the extracellular ATD and ABD; the membrane TMD with transmembrane segments M1, M3, M4, and the M2 pore loop forming the ion channel; and the intracellular CTD. Graph showing the amino acid alignment between hGluN2A (NP_001127879.1) and each of the zebrafish GluN2A paralogs zGluN2Aa and zGluN2Ab (XP_021329529.1, XP_009304490.1). Values indicate percent homology derived from ClustalW alignment between the indicated subunits using a sliding average of 25 amino acid segments. Above the graph is the linear representation of the hGluN2A subunit. The colors correspond to the cartoon, with the ATD shown in blue, the ABD in green, the TMD in yellow, and the CTD in gray. The amino acid numbers that make up each domain are indicated. B, C, Representative current recordings in HEK293T cells expressing zGluN1a/zGluN2Aa or zGluN1b/zGluN2Bb receptors (B) or hGluN1/hGluN2A or hGluN1/GluN2B receptors (C), with responses evoked by a brief (10–15 ms) application of 1 mM glutamate in the continuous presence of 100 µM glycine. The responses normalized to the peak amplitude are shown on the right (red lines indicate the double exponential function fit to the data: τ_w_ =36.2 ms for zGluN1a/zGluN2Aa and τ_w_ = 118.9 ms for zGluN1b/zGluN2Bb; τ_w_ = 58.8 ms for hGluN1/hGluN2A and τ_w_ = 510 ms for hGluN1/hGluN2B).

The structural similarities between human GluN2A (hGluN2A, encoded by *GRIN2A*) and the zebrafish paralogs zGluN2Aa and zGluN2Ab (encoded by *grin2Aa* and *grin2Ab*, respectively) were analyzed. zGluN2Aa and zGluN2Ab share high protein sequence homology, particularly in the transmembrane domain (TMD) and the S1 and S2 segments of the agonist-binding domain (ABD) (85–100%) (Fig. 1A). The channel-proximal region of the amino-terminal domain (ATD, lower lobe) shows intermediate similarity (55–93%), while the distal ATD (upper lobe) and the C-terminal domain (CTD) display lower homology (25–85% and 10–80%, respectively). Comparisons with hGluN2A reveal similarly high conservation in the ABD and TMD. Despite lower sequence identity in the CTD, both zebrafish subunits retain conserved motifs involved in receptor trafficking and synaptic targeting (Won et al., 2017). Notably, the last four amino acids—ESVD in zGluN2Aa (1457–1460) and zGluN2Ab (1442– 1445)—form a canonical PDZ-binding motif critical for interactions with membrane-associated guanylate kinases (MAGUKs) (Kim and Sheng, 2004).

The functional properties of rat NMDARs are governed by their subunit composition (Monyer et al., 1994), which also influences the kinetics of synaptic transmission. Receptors enriched in GluN2B subunits, characteristic of early postnatal hippocampal neurons, display slow deactivation, whereas those enriched in GluN2A subunits, characteristic of adult hippocampal neurons, show faster deactivation kinetics (Flint et al., 1997) (Gray et al., 2011). Triheteromeric GluN1/GluN2A/GluN2B receptors display intermediate deactivation rates compared to GluN1/GluN2A and GluN1/GluN2B diheteromers (Hansen et al., 2014). To test whether zebrafish NMDARs show similar subunit-dependent deactivation kinetics, we co-expressed zebrafish GluN2A and GluN2B subunits (zGluN2Aa, zGluN2Ab, zGluN2Ba, zGluN2Bb) with zebrafish GluN1a or GluN1b (zGluN1a, zGluN1b) subunits in HEK293T cells. Note that the two zGluN1 paralogs we used contained the alternatively spliced exon 5 (Cox et al., 2005). Using whole-cell patch-clamp recordings, we assessed the current density of glutamate-evoked responses as well as the deactivation time course of NMDAR responses following brief application. In the continuous presence of 1 mM glycine, 1 mM glutamate elicited current densities ranging from 0.2 to 0.6 nA/pF across most receptor combinations, indicative of functional surface expression. However, receptors formed with zGluN2Ba (XP_009297288.1, 1353 aa) generated minimal currents (2.7 ± 1.1 pA/pF; n = 6), preventing further characterization of this subunit.

The time course of NMDAR response decay following a brief application of glutamate (1 mM) in the presence of saturating glycine is commonly used to approximate the kinetics of synaptic NMDAR currents (Lester et al., 1990) (Fig. 1B). Zebrafish NMDARs containing zGluN2Aa or zGluN2Ab subunits showed fast deactivation kinetics, regardless of whether they were co-expressed with zGluN1a or zGluN1b. The weighted time constants (τ_w_) were as follows: 42.9 ± 4.8 ms (*n* = 5) for zGluN1a/zGluN2Aa, 30.9 ± 1.6 ms (*n* = 11) for zGluN1b/zGluN2Aa, 32.5 ± 2.6 ms (*n* = 5) for zGluN1a/zGluN2Ab, and 38.1 ± 3.5 ms (*n* = 5) for zGluN1b/zGluN2Ab. In contrast, receptors containing the zGluN2Bb subunit exhibited slower deactivation: 108.6 ± 23.6 ms (*n* = 5) for zGluN1a/zGluN2Bb and 146.8 ± 42.5 ms (*n* = 5) for zGluN1b/zGluN2Bb. For comparison, human NMDARs deactivated with τ_w_ of 60.3 ± 6.3 ms (*n* = 6) for hGluN1/hGluN2A and 604 ± 40 ms (*n* = 6) for hGluN1/hGluN2B. Although zebrafish NMDARs generally deactivate faster than their mammalian counterparts, the subunit-dependent pattern is conserved: GluN2A-containing receptors deactivate rapidly, while GluN2B-containing receptors show slower kinetics (Hansen et al., 2021).

### Expression of zebrafish *grin2A* paralogs in early development

Previous studies have shown that zebrafish NMDAR subunit mRNAs are differentially expressed in the developing nervous system (Cox et al., 2005; Zoodsma et al., 2020; Zoodsma et al., 2022). Our *in-situ* hybridization (ISH) analysis revealed no or minimal expression of *grin2Aa* or *grin2Ab* mRNA in the brain, retina (Fig. 2A, B), or spinal cord (data not shown) at 1 dpf. By 3 dpf, both transcripts were detectable in the nervous system: *grin2Aa* was localized predominantly to the optic tectum, while *grin2Ab* showed broader expression throughout the brain and retina (Fig. 2C, D, I). At 5 dpf, *grin2Aa* expression became more pronounced in the telencephalon, tectal proliferation zone, cerebellum, hindbrain, and retina. *Grin2Ab* remained strongly expressed in the optic tectum and retina, but at lower levels in other brain regions (Fig. 2E, F, I). While *grin2Aa* and *grin2Ab* displayed overlapping expression patterns—especially in the retina—*grin2Aa* generally exhibited higher intensity labeling in the brain. In the spinal cord, both genes showed limited expression at all time points analyzed, with signal levels comparable to negative controls using sense probes (Fig. 2G, H).

**Figure 2.**
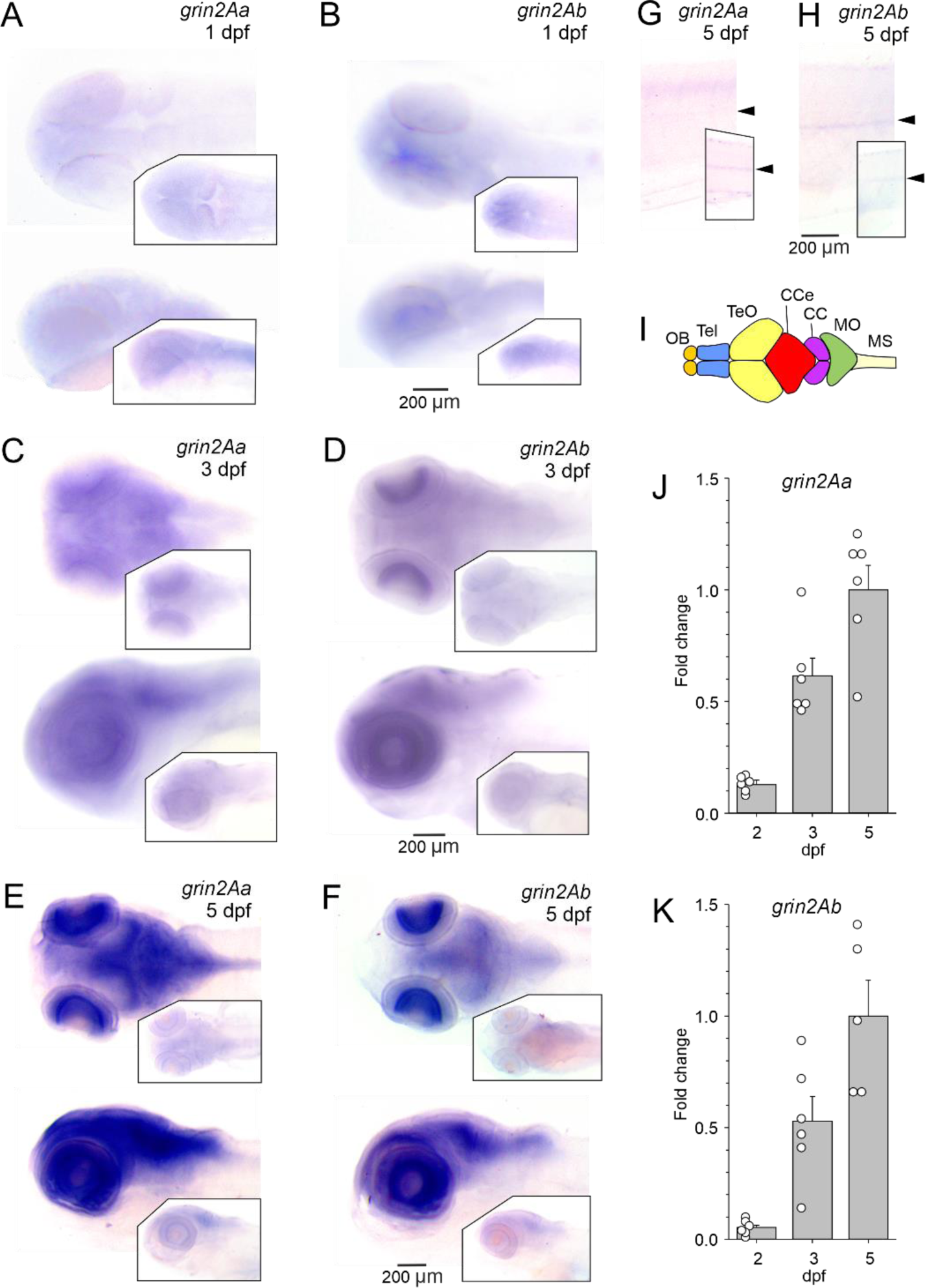
Expression of grin2Aa and grin2Ab in the zebrafish nervous system. Whole-mount ISH of *grin2Aa* and *grin2Ab* at 1 dpf (A, B), 3 dpf (C, D), and 5 dpf (E, F) showing dorsal (top) and lateral (bottom) views. A, C, E, Insets, sense probes of *grin2Aa*. B, D, F, Insets, sense probes of *grin2Ab*. G, H, Lateral view of the trunk at 5 dpf for *grin2Aa* (G) and *grin2Ab* (H). The position of the spinal cord is indicated by arrows. Insets, sense probes of *grin2Aa* and *grin2Ab*. I, Simplified representation of the zebrafish central nervous system and its major structures. Adapted from (Wullimann et al., 1996); CC: crista cerebellaris; CCe: Corpus cerebelli; MO: medulla oblongata; MS: medulla spinalis; OB: olfactory bulb; Tel: telencephalon; TeO: tectum opticum. J, K, Relative expression levels of *grin2Aa* and *grin2Ab* mRNA at 2 dpf (*n* = 6; 6), 3 dpf (*n* = 6; 6), and 5 dpf (*n* = 6; 5) in the head of zebrafish larvae analyzed using RT-qPCR. Results are normalized to the corresponding gene expression at 5 dpf.

The ISH experiments suggest that *grin2Aa* and *grin2Ab* expression begins after 1 dpf and increases progressively during early larval development. To validate and quantify this trend, we performed RT-qPCR on mRNA isolated from whole larval heads. Consistent with the ISH results, RT-qPCR revealed low expression levels of *grin2Aa* and *grin2Ab* at 2 dpf, followed by a gradual increase through 5 dpf (Fig. 2J, K). Similarly, the expression levels of other NMDAR subunit paralogs—*grin1a*, *grin1b*, *grin2Ba*, and *grin2Bb*—also increased during the 2–5 dpf period (Fig. S1), mirroring the developmental expression patterns reported in the central nervous system of higher vertebrates (Tovar and Westbrook, 1999).

### Generation of *grin2Ab* knockout zebrafish

In recent years, zebrafish lines lacking zGluN1 (*grin1a*^−/−^ and *grin1b*^−/−^) and zGluN2B (*grin2Ba*^−/−^ and *grin2Bb*^−/−^) subunits have been developed to study the role of NMDARs in early brain development and behavior (Zoodsma et al., 2020; Zoodsma et al., 2022; Thyme et al., 2019; Griffin et al., 2021). To investigate the role of GluN2A in zebrafish, we utilized two loss-of-function mutant strains: *grin2Aa* and *grin2Ab.* The *grin2Aa* mutant harbors a point mutation (T>A) at chr3:27,207,119 (GRCz11) in exon 5 (ENSDART00000142141.5), converting the leucine codon (TTG) to a premature stop codon (TAG), resulting in a truncated protein of 344 amino acids containing most of the ATD (zGluN2Aa-L345Ter) (Dooley et al., 2013; Kettleborough et al., 2013) (Fig. 3A, B). The *grin2Ab* mutant was generated via CRISPR/Cas9, introducing a 16-bp deletion in exon 4 (chr1:8,810,711–8,810,726; ENSDART00000103652.5), leading to a frameshift and premature stop codon (zGluN2Ab-E232GfsTer14), truncating the protein at 244 amino acids, preserving approximately half of the ATD (Fig. 3A-C). Additionally, a *grin2Bb* mutant was created using CRISPR/Cas9 by deleting an adenosine at chr1:45,449,325 in exon 4 (ENSDART00000143664.3), producing a frameshift and stop codon (zGluN2Bb-I331SfsTer21), resulting in a 350-amino-acid truncated protein containing most of the ATD, similar to the *grin2Aa* mutant (Fig. 3A-C). Because all three mutations introduce premature stop codons upstream of essential domains for co-assembly with zGluN1 and receptor trafficking, the resulting proteins are expected to be non-functional.

**Figure 3.**
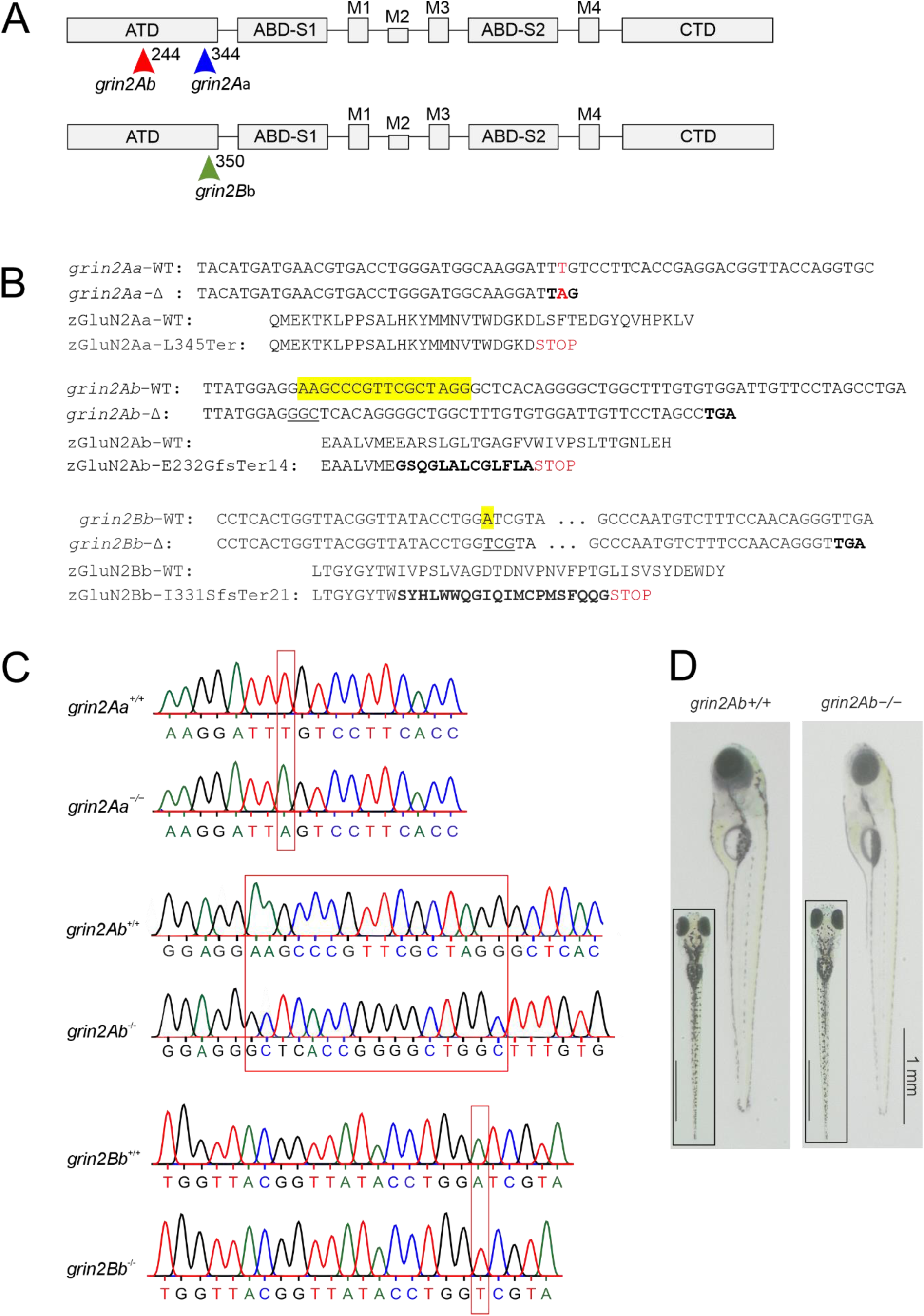
Locations of zGluN2 mutations resulting in a premature stop codon and protein truncation. A, Schematic representation of the domain structure of a GluN2 subunit. It consists of four domains: the extracellular ATD and S1 and S2 lobes of the ABD; the membrane TMD with transmembrane segments M1, M3, M4, and M2 pore loop forming the ion channel; and the intracellular CTD. In the ATD, arrows indicate the position of *grin2Aa* mutation (sa14573), and gRNA target sites for *grin2Ab* and *grin2Bb*. B, The nucleotide and amino acid sequence alignment of the portion of the ATD of zGluN2Aa wild-type (*grin2Aa*-WT; zGluN2Aa-WT)) and the mutant strain (*grin2Aa*-Δ; zGluN2Aa-L345Ter) (*top*), zGluN2Ab wild-type (*grin2Ab*-WT; zGluN2Ab-WT) and the mutant strain (*grin2Ab*-Δ; zGluN2Ab-E232GfsTer14) (*middle*), and zGluN2Bb wild-type (*grin2Bb*-WT; zGluN2Bb-WT) and the mutant strain (*grin2Bb*-Δ; zGluN2Bb-I331SfsTer21) (*bottom*). C, Chromatograms of Sanger sequencing of PCR amplicons of genomic DNA from *grin2Aa^+/+^* and *grin2Aa^−/−^* (*top*), *grin2Ab^+/+^* and *grin2Ab^−/−^* (*middle*), and *grin2Bb^+/+^* and *grin2Bb^−/−^* (*bottom*) D, Lateral and dorsal (inset) images of representative 6 dpf wild type (*grin2Ab*^+/+^) and *grin2Ab^−/−^* larvae.

Heterozygous *grin2Aa*^+/−^ and *grin2Ab*^+/−^ embryos and larvae were viable, raised to adulthood, genotyped, and intercrossed to generate homozygous mutants *grin2Aa*^−/−^, *grin2Ab*^−/−^. Homozygous double mutants *grin2Aa*^−/−^ *grin2Ab*^−/−^ lacking all GluN2A subunits (*grin2A*^−/−^) were obtained by crossing heterozygous double mutants *grin2Aa*^+/−^ *grin2Ab*^−/−^, *grin2Aa*^−/−^ *grin2Ab*^+/−^, or homozygous double mutants *grin2Aa^−/−^ grin2Ab^−/−^*. Expected and observed recovery rates of the different genotypes at 6 dpf are summarized in Table S2. Similar to *grin2Aa*^−/−^ and *grin2Ab*^−/−^, *grin2A*^−/−^ larvae appear morphologically normal at 6 dpf (Fig. 3D) were viable and survived to adulthood. Analysis of fish standard length (from snout to the tail peduncle) showed that *grin2Aa^−/−^*, *grin2Ab^−/−^*, and *grin2A^−/−^* fish reached similar sizes at 56, 70, 84 and 98 dpf as their wild-type siblings (*grin2Aa^+/+^ grin2Ab^+/+^*) (One-way ANOVA showed no differences in the standard length among the genotypes at 56 (*p* = 0.654), 70 (*p* = 0.999), 84 (*p* = 0.784), and 98 (*p* = 0.395) dpf) (Fig. S2).

These findings are consistent with previous reports in mice, where GluN2A knockout animals are viable and develop normally (Sakimura et al., 1995), in contrast to GluN2B knockout mice, which die shortly after birth (Kutsuwada et al., 1996). Similarly, our independently generated *grin2Bb* mutant zebrafish were recovered at Mendelian ratios and survived to adulthood, as reported previously for the zGluN2Bb-V699EfsTer3 mutant line (Zoodsma et al., 2022).

### Zebrafish lacking *grin2A* exhibit increased locomotion and altered thigmotaxis

To investigate the role of zGluN2A and zGluN2B subunits in zebrafish behavior, we employed a swim behavior assay analogous to the open field test, commonly used in behavioral neuroscience to assess novelty response, exploration, and anxiety (Walsh and Cummins, 1976). In this test, larvae were placed in an open arena, and their spatio-temporal locomotion patterns were analyzed. Swimming behavior was assessed in zebrafish larvae at 6 dpf. Figure 4A, B shows the recording setup and a representative tracking plot of larval movement. Larval activity consisted of short swimming bouts interspersed with rest periods (Fig. 4C). To quantify locomotion, we analyzed three parameters over 10-minute intervals: mean travel distance (MTD), mean bout distance (MBD), and mean bout frequency (MBF).

**Figure 4.**
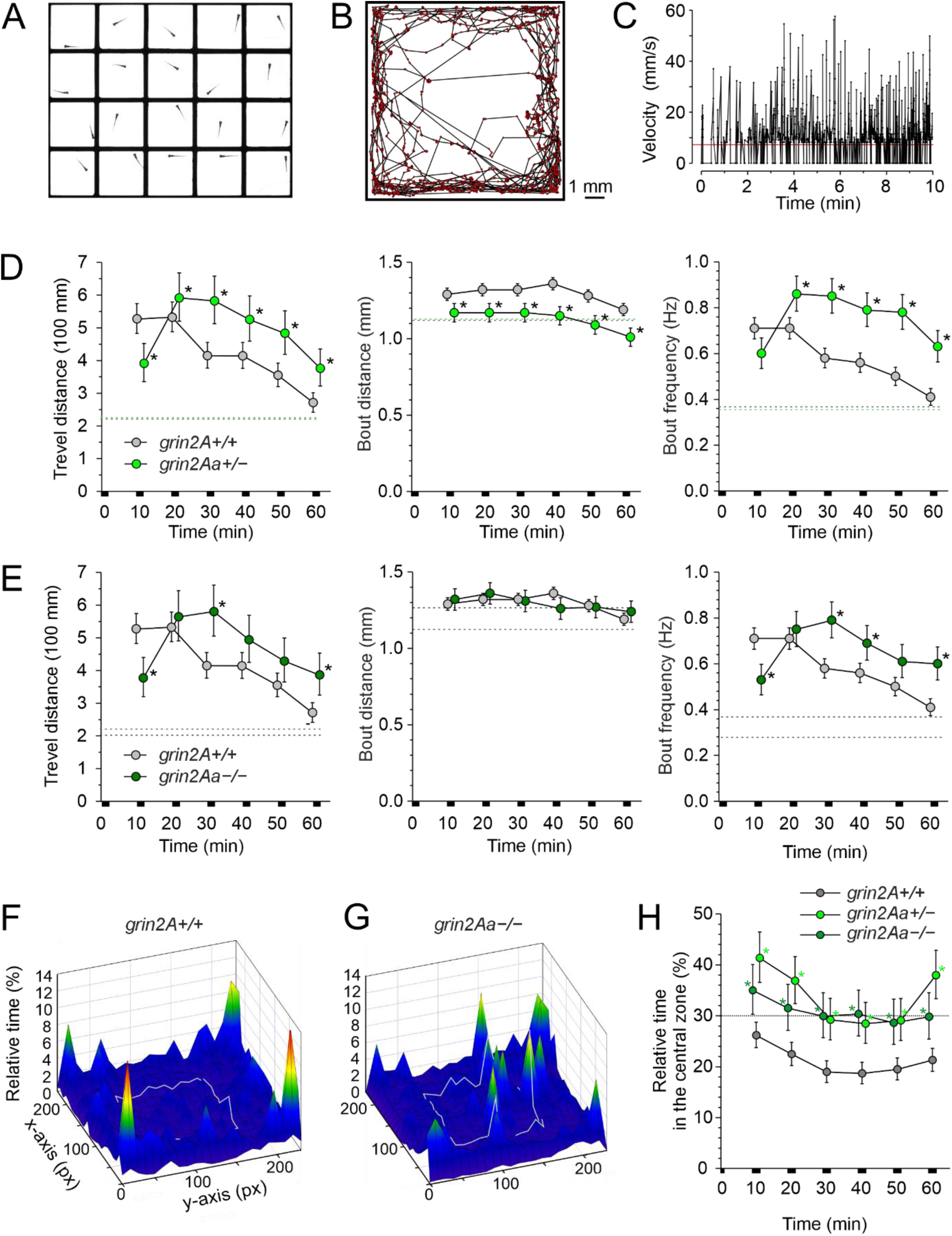
Effects of *grin2Aa* deletion on larval swimming behavior and thigmotaxis. A, Photograph of a multi-well chamber used to simultaneously assess the spontaneous swimming behavior of 20 larvae (6 dpf). Larvae in the experiment were progeny of *grin2Aa^+/−^* intercrosses with the genotype determined after the behavioral experiment. B, The graph shows the sample trajectory of a *grin2Aa^+/+^* larva as assessed during the first 10 min interval. Data points were acquired at 50 Hz. C, The graph shows the velocity of a *grin2A^+/+^* larva assessed at 20 ms intervals. The red line indicates the velocity threshold for locomotor activity detection (see Methods). The MTD, MBD, and MBF of *grin2Aa^+/−^* and *grin2A^+/+^* larvae (D) and *grin2Aa^−/−^* and *grin2A^+/+^* larvae (E) are compared. Dashed lines correspond to mean values of the measured parameters at steady state, 2–3 h after the placement of larvae into the experimental chamber. Representative heat maps for the relative time spent by *grin2A^+/+^* (F) and *grin2Aa^−/−^* (G) larvae in 0.454 x 0.454 mm area bins (10 x 10 pixels) over 20 min of recording (30–50 min), indicated by color coding. The central zone (5.5 x 5.5 mm) used to assess thigmotaxis is outlined by a gray line. H, Comparison of thigmotaxis measured as the relative time spent by *grin2A^+/+^*, *grin2Aa^+/−^*, and *grin2Aa^−/−^* larvae in the central zone. The analysis was performed on the same larvae used to study locomotion. The dotted line indicates the level of randomness (30%). Data are expressed as mean ± 95% confidence interval of travel distance, bout distance, and bout frequency (D, E), and the relative time spent by larvae in the central zone (H) in episodes lasting 10 min; * indicates significant differences in genotype as assessed by ANOVA followed by LSD *post-hoc* test (*grin2A^+/+^*: *n* = 582; *grin2Aa^+/−^*: *n* = 429; *grin2Aa^−/−^*: *n* = 216). *grin2Aa^+/−^ versus grin2A^+/+^*: travel distance genotype: *p* < 0.001, time: *p* < 0.001; bout distance genotype: *p* < 0.001, time: *p* < 0.001; bout frequency: genotype: *p* < 0.001, time: *p* < 0.001; and relative time larvae spent in the central zone genotype: *p* < 0.001, time: *p* < 0.001. *grin2Aa^−/−^ versus grin2A^+/+^*: travel distance genotype: *p* = 0.0056, time: *p* < 0.001; bout distance genotype: *p* = 0.81, time *p* = 0.013; bout frequency: genotype: *p* = 0.001, time: *p* < 0.001; and relative time spent by larvae in the central zone genotype: *p* < 0.001, time: *p* = 0.013.

Figure 4D shows locomotor analysis of wild-type control larvae. The MTD was stable during the first (0–10 min) and second (10–20 min) intervals (0.527 m and 0.532 m, respectively) and gradually declined over subsequent intervals. The MBD during the first interval was 1.29 mm and remained relatively constant throughout the session. The MBF was 0.71 Hz in the first interval and declined similarly to MTD. No significant differences in MTD, MBD, or MBF were observed between wild-type sibling larvae from *grin2Aa* or *grin2Ab* mutant crosses; therefore, the data were pooled and presented as *grin2A^+/+^*.

The MTD of *grin2Aa^+/−^* larvae during the first 10 min interval was lower than in *grin2A^+/+^* larvae, but increased 1.5-fold during the second interval (Fig. 4D). From 20–60 min, MTD remained significantly higher in *grin2Aa^+/−^* larvae compared to *grin2A^+/+^*, though both genotypes showed a similar decline over time. The MBD in *grin2Aa^+/−^* larvae was slightly but significantly lower than in *grin2A^+/+^*during the first interval (1.17 mm) and remained stable throughout. The MBF followed a pattern similar to MTD, increasing 1.4-fold in the second interval and remaining elevated compared to *grin2A^+/+^*. *grin2Aa^−/−^*larvae exhibited a comparable profile to *grin2Aa^+/−^* larvae, with a milder increase in MTD and MBF; MBD was not significantly different from controls.

To assess steady-state locomotion in *grin2A^+/+^*, *grin2Ab^+/−^*, and *grin2Ab^−/−^* larvae, we conducted additional experiments in which swimming parameters were measured 2–3 hours after larvae were placed into the experimental wells. At this time point, none of the three parameters differed significantly between genotypes, nor did they show any change across the six 10-minute intervals analyzed (Fig. S4). Therefore, the data were pooled and presented as dotted lines in the corresponding panels of Figure 4. Together, the data suggest that both heterozygous and homozygous *grin2Aa* deletion results in transient hyperlocomotion in response to a novel environment.

Locomotor activity was also used to assess thigmotaxis by measuring the relative time larvae spent in the central zone of the well. Thigmotaxis, an established index of anxiety in zebrafish and other species (Schnorr et al., 2012), reflects the tendency to avoid open areas and stay near boundaries. *grin2A^+/+^*larvae explored the entire well but spent only 18.7% to 26.2% of each 10-minute interval in the central zone—less than the 30% chance level based on zone area (Fig. 4F, H; dotted line). In contrast, *grin2Aa^+/−^* and *grin2Aa^−/−^*larvae spent 28.5% to 41.4% of each interval in the center, significantly more than *grin2A^+/+^* controls (Fig. 4G, H). These results suggest that wild-type larvae exhibit mild anxiety-like behavior in a novel environment, while deletion of one or both *grin2Aa* alleles reduces anxiety.

We next assessed whether deletion of the *grin2Ab* paralog alters zebrafish behavior in a novel environment. In heterozygous *grin2Ab^+/−^*larvae, swimming parameters—including MTD, MBD, and MBF—were largely comparable to *grin2A^+/+^* controls, with the exception of a modest increase in MTD during the 20–30 min interval (Fig. 5A). In contrast, homozygous *grin2Ab^−/−^* larvae exhibited significantly elevated MTD and MBF across all time intervals from 10 to 60 min (Fig. 5B), though not under steady-state conditions 2–3 hours post-transfer (dotted lines in Fig. 5A, B). MBD values in *grin2Ab^−/−^* larvae remained similar to those in *grin2A^+/+^* larvae at all time points. To further explore the functional role of *grin2A*, we generated double mutant zebrafish lacking both *grin2Aa* and *grin2Ab* (*grin2A^−/−^*). Behavioral analysis revealed that *grin2A^−/−^* larvae exhibited significantly increased MTD and MBF during 10-minute intervals compared to *grin2A^+/+^*controls, while MBD remained unchanged—mirroring the phenotypes observed in *grin2Aa^−/−^* and *grin2Ab^−/−^* larvae (Fig. 4E; Fig. 5A-C).

**Figure 5.**
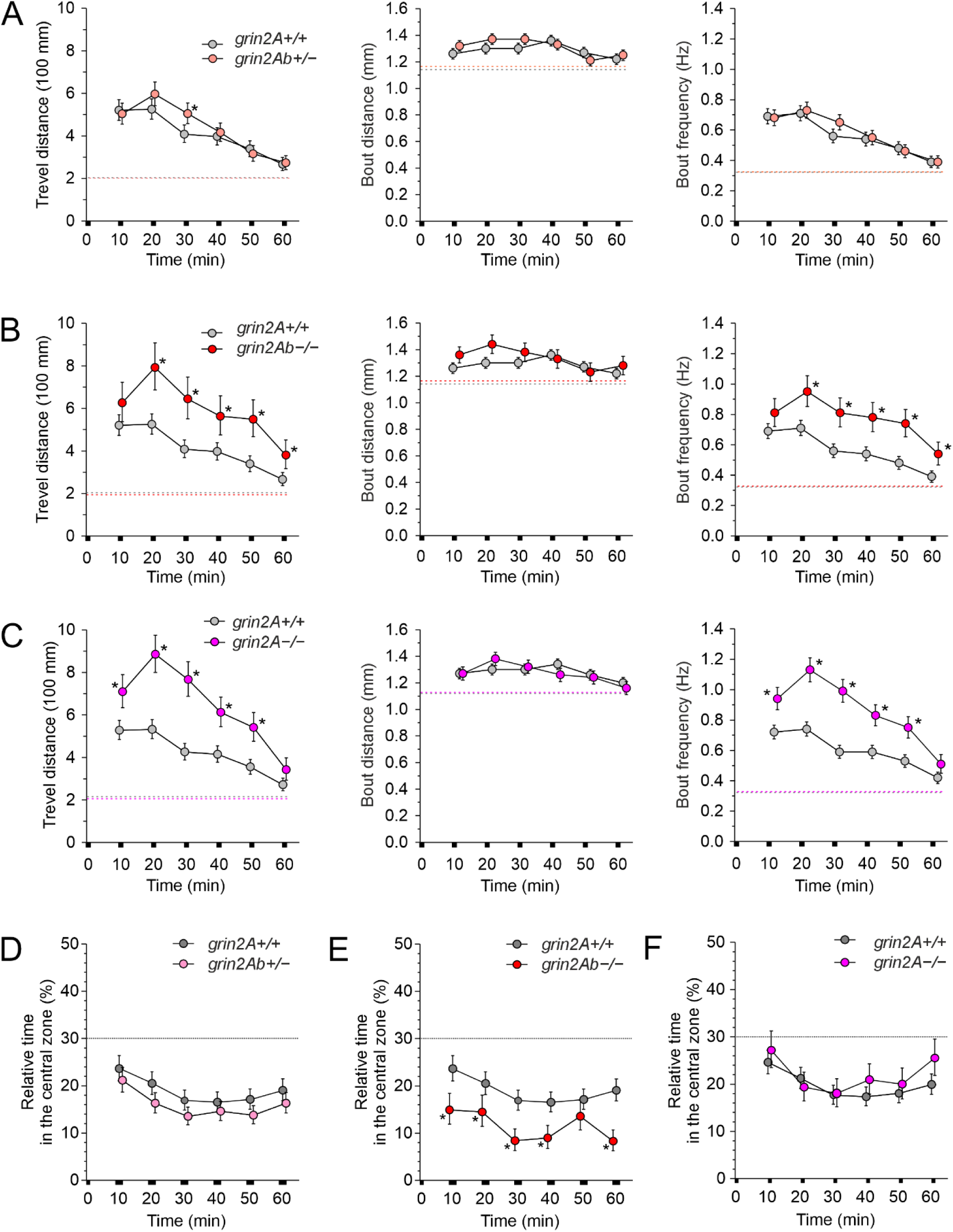
Effects of *grin2Ab* deletion on larval swimming behavior and thigmotaxis. Comparison of MTD, MBD, and MBF of *grin2Ab^+/−^* (A), *grin2Ab^−/−^* (B), and *grin2A^−/^*^−^ (C) larvae compared to *grin2A^+/+^* larvae (6 dpf). Mean steady-state values of the locomotion parameters examined 2–3 h after the placement of larvae in the experimental wells are shown as dashed lines. D, Comparison of thigmotaxis measured as the relative time spent in the central zone by *grin2Ab^+/−^*, *grin2Ab^−/−^*, and *grin2A^−/^*^−^ larvae compared to *grin2A^+/+^* larvae. The analysis of thigmotaxis was performed on the same larvae used to study locomotion. The dotted line indicates the level of randomness (30%). Data are expressed as mean ± 95% confidence interval of travel distance, bout distance, and bout frequency (A–C), and the relative time spent by larvae in the central zone (D) measured in 10 min intervals; * indicates significant differences in the genotype as assessed by ANOVA followed by LSD *post-hoc* test (*grin2A^+/+^*: *n* = 584; *grin2Ab^+/−^*: *n* = 542; *grin2Ab^−/−^*: *n* = 197; *grin2A^−/−^*: *n* = 270). *grin2Ab^+/−^ versus grin2A^+/+^*: travel distance genotype: *p* = 0.26, time: *p* < 0.001; bout distance genotype: *p* = 0.062, time: *p* < 0.001; bout frequency genotype: *p* = 0.55, time: *p* < 0.001; the relative time larvae spent in the central zone genotype: *p* = 0.001, time: *p* < 0.001. *grin2Ab^−/−^ versus grin2A^+/+^*: travel distance genotype: *p* < 0.001, time: *p* < 0.001; bout distance genotype: *p* = 0.051, time: *p* = 0.003; bout frequency genotype: *p* = 0.001, time: *p* < 0.001; the relative time larvae spent in the central zone genotype: *p* < 0.001, time: *p* = 0.001. *grin2A^−/−^ versus grin2A^+/+^*: travel distance genotype: *p* < 0.001, time: *p* < 0.001; bout distance genotype: *p* = 0.50, time: *p* < 0.001; bout frequency genotype: *p* < 0.001, time: *p* < 0.001; and the relative time larvae spent in the central zone genotype: *p* = 0.10, time: *p* < 0.001.

Thigmotaxis analysis showed that *grin2Ab^+/−^* larvae avoided the center zone similarly to *grin2A^+/+^* larvae, whereas *grin2Ab^−/−^* larvae spent significantly less time in the center (Fig. 5D,E), suggesting heightened anxiety-like behavior. However, unlike *grin2Aa^−/−^*larvae, *grin2A^−/−^* larvae did not differ significantly from controls in the time spent in the central zone of the well, suggesting no major alterations in thigmotaxis (Fig. 5F).

Together, the behavioral findings indicate that while deletion of either *grin2A* paralog increases larval locomotion, only *grin2Aa^+/−^*larvae exhibit haploinsufficiency, whereas *grin2Ab^+/−^* larvae are haplosufficient. Moreover, thigmotaxis analysis suggests opposing roles for the two paralogs in anxiety regulation: *grin2Aa* deletion appears anxiolytic, while *grin2Ab* deletion is anxiogenic. These effects may counterbalance each other in *grin2A^−/−^* larvae, resulting in no net change in anxiety-related behavior.

### Deletion of *grin2Bb* does not impact zebrafish locomotor activity and thigmotaxis

To further examine the subunit-specific contributions of NMDARs to zebrafish larval behavior, we assessed the effects of mono- and bi-allelic deletion of the *grin2Bb* gene using the same locomotion assay applied to *grin2A* mutants. Analysis of locomotor activity and thigmotaxis in heterozygous *grin2Bb^+/−^* larvae revealed no significant differences in MTD, MBD, or MBF compared to wild-type siblings (*grin2B^+/+^*) across all time intervals (Fig. 6A). Similarly, homozygous *grin2Bb^−/−^* larvae exhibited locomotion parameters indistinguishable from *grin2B^+/+^* and *grin2Bb^+/−^*controls (Fig. 6B). Thigmotaxis, measured as the relative time spent in the central zone, also did not differ significantly between genotypes (Fig. 6C). Collectively, these results suggest that deletion of *grin2Bb* does not alter swimming behavior or anxiety-like responses in zebrafish larvae.

**Figure 6.**
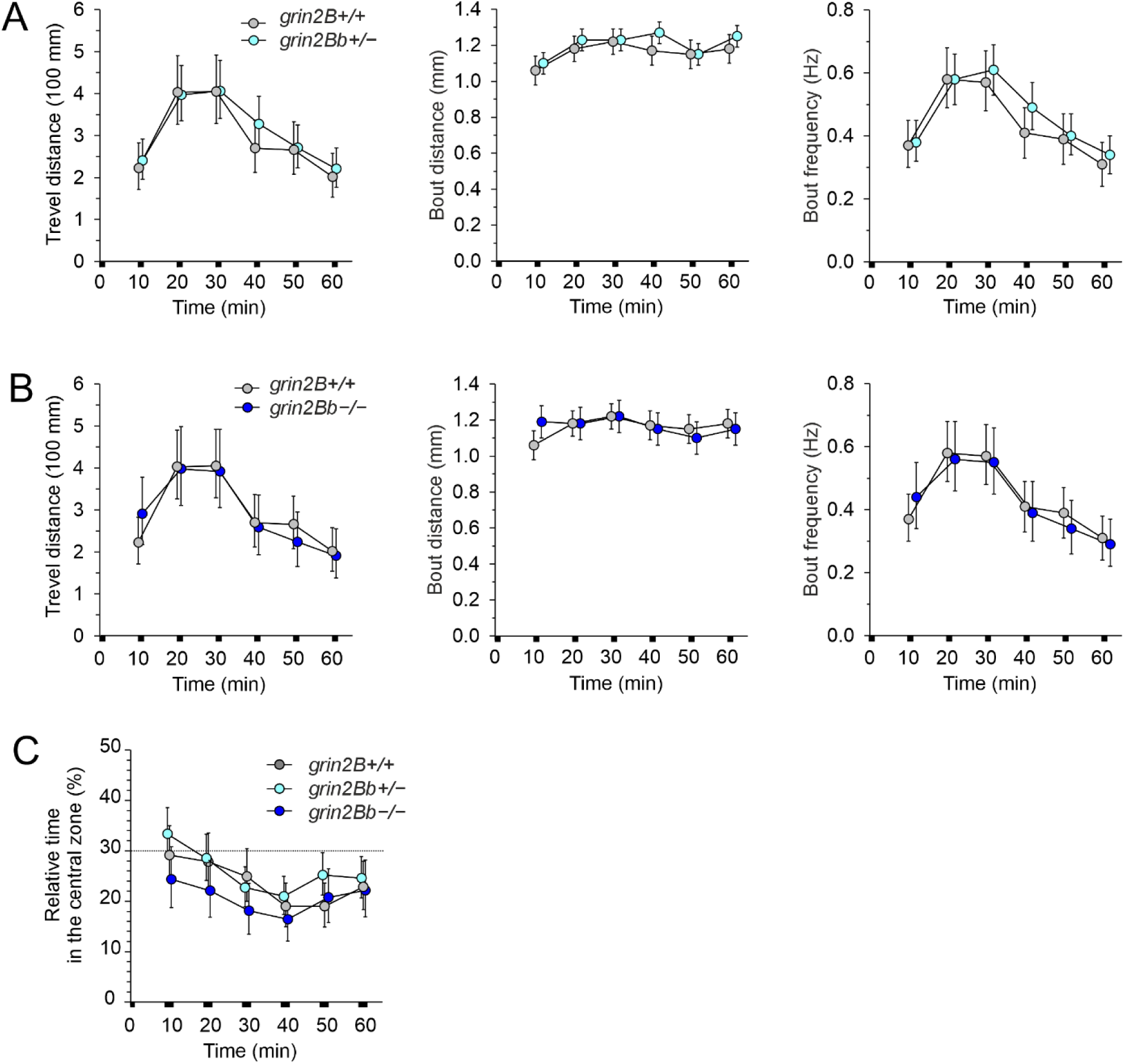
No effect of *grin2Bb* deletion on larval locomotor activity and thigmotaxis. Comparison of the MTD, MBD, and MBF assessed in *grin2Bb^+/−^* (A), and *grin2Bb^−/^*^−^ (B) larvae compared to *grin2B^+/+^* larvae (6 dpf). C, Thigmotaxis measured as the relative time spent in the central zone by *grin2Bb^+/−^* and *grin2Bb^−/−^* larvae compared to *grin2B^+/+^* larvae. The analysis of thigmotaxis was performed on the same larvae used to study locomotion. The dotted line indicates the level of randomness (30%). Data are expressed as mean ± 95% confidence interval of travel distance, bout distance and bout frequency (A–B), and the relative time spent by larvae in the central zone (C); data were analyzed by ANOVA followed by *post-hoc* LSD tests (*grin2B^+/+^*: *n* = 198; *grin2Bb^+/−^*: *n* = 288; *grin2Bb^−/−^*: *n* = 144). *grin2Bb^+/−^ versus grin2B^+/+^*: travel distance genotype: *p* = 0.18, time: *p* < 0.001; bout distance genotype: *p* = 0.37, time: *p* < 0.003; bout frequency genotype: *p* = 0.13, time: *p* < 0.002; the relative time larvae spent in the central zone genotype: *p* = 0.23, time: *p* < 0.002. *grin2Bb^−/−^ versus grin2B^+/+^*: travel distance genotype: *p* = 0.97, time: *p* < 0.001; bout distance genotype: *p* = 0.53, time: *p* = 0.35; bout frequency genotype: *p* = 0.83, time: *p* < 0.001; the relative time larvae spent in the central zone genotype: *p* = 0.11, time: *p* = 0.23.

### Compensatory changes in NMDAR subunit expression in mutant *grin2Aa* and *grin2Ab* fish

Organisms exhibit genetic robustness—the capacity to buffer the phenotypic impact of mutations (de Visser et al., 2003). To explore whether nonsense mutations in *grin2Aa* and *grin2Ab* trigger transcriptional downregulation of mutant mRNA or compensatory upregulation of functionally related genes, we next examined potential mechanisms of molecular compensation (Rouf et al., 2023).

RT-qPCR was used to quantify mRNA expression of six NMDAR subunits in 6 dpf larvae (Fig. 7A-C). In *grin2Aa^−/−^* larvae, *grin2Aa* mRNA levels were comparable to *grin2A^+/+^* controls (Fig. 7B). In contrast, *grin2Ab* mRNA was significantly downregulated in *grin2Ab^−/−^* larvae, accompanied by a 1.4-fold upregulation of *grin2Aa* mRNA. In *grin2A^−/−^* double knockouts, *grin2Ab* transcript levels were also reduced, but no additional changes were observed (Fig. 7B). No significant differences were found in the expression of *grin1* or *grin2B* paralogs across genotypes (Fig. 7A, C). These results suggest that only *grin2Ab^−/−^* larvae exhibit compensatory transcriptional regulation, potentially through nonsense-mediated mRNA decay (NMD), which degrades aberrant transcripts and may trigger upregulation of related genes (Nagy and Maquat, 1998).

**Figure 7.**
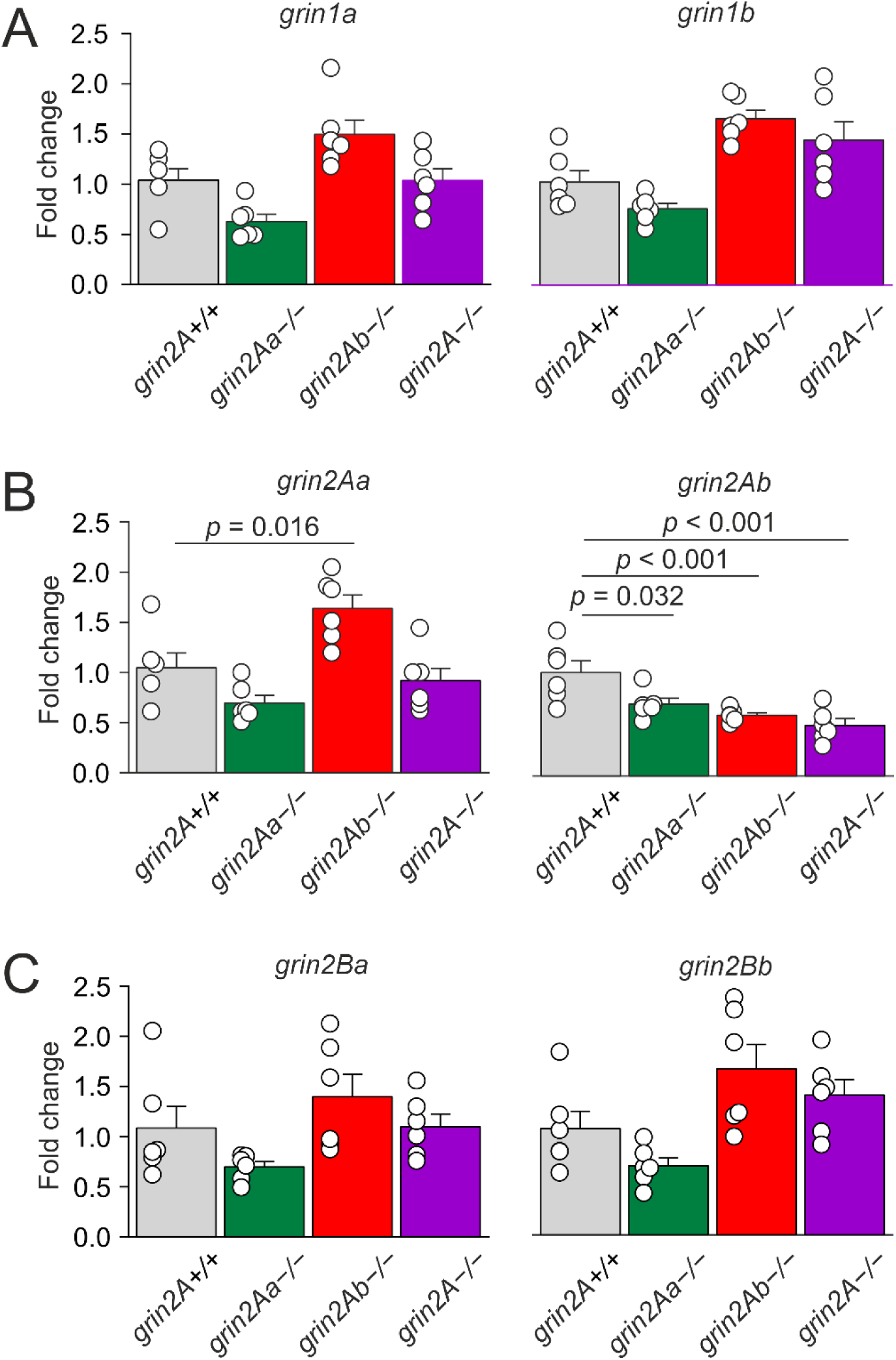
Effects of *grin2Aa* or *grin2Ab* deletion on *grin* mRNA expression in zebrafish larvae. Relative expression levels of *grin1* (A), *grin2A* (B) and *grin2B* (C) mRNA in heads of zebrafish larvae (6 dpf) for *grin2A^+/+^*, *grin2Aa^−/−^*, *grin2Ab^−/−^*, and *grin2A^−/−^* fish were analyzed by RT-qPCR. One-way ANOVA was used to assess the differences in the relative expression of each gene, followed by multiple comparisons *versus* the corresponding gene in the grin2A+/+ larvae (Dunnett’s method), with significance as indicated. *grin1a: p* = 0.028; *grin1b*: *p* = 0.010; *grin2Aa*: *p* < 0.001; *grin2Ab*: *p* < 0.001; *grin2Ba*: *p* = 0.030; *grin2Bb*: *p* = 0.011; *n* = 6 for each genotype: *grin2A+/+, grin2Aa−/−, grin2Ab−/− and grin2A−/−,* for each analysis.

We also examined the impact of *grin2A* paralog deletion on the expression of *grin1*, *grin2A*, and *grin2B* mRNAs in the nervous tissue of adult *grin2Aa^−/−^*, *grin2Ab^−/−^* and *grin2A^+/+^* zebrafish. Analyses were conducted in the optic tectum, a clearly defined brain region with high NMDAR expression. No significant changes in NMDAR subunit mRNA levels were observed in adult mutants, except for a significant downregulation of *grin2Ab* mRNA in *grin2Ab^−/−^* fish (Fig. S3).

### Protein expression in the nervous tissue of WT and mutant fish

To investigate protein expression changes resulting from the loss of GluN2Aa or GluN2Ab, we performed mass spectrometry-based proteomic analysis on the optic tectum from adult zebrafish. Tissue samples were collected from five *grin2A^+/+^*, four *grin2Aa*^−/−^, and four *grin2Ab*^−/−^ fish. After applying filtering criteria detailed in the Methods section, a total of 6,615 proteins were identified for downstream analysis.

Bi-allelic deletion of *grin2Aa* significantly altered the expression of 55 proteins in adult zebrafish (defined as <0.5-fold or >2-fold change relative to *grin2A⁺/⁺* controls; Fig. 8A, Table S3). The most upregulated proteins included an anion exchange protein (D1FTM8; 25-fold) and several myosin family members (P13104, Q6IQX1, O93409, B6IDE1, A0A8M9PJW6, F1R6C7, E9QF07), with increases ranging from 5-to 8.9-fold. Downregulated proteins included calbindin 2 (Q6PC56; 0.36-fold) and neuregulin 2 (E7FGA4; 0.36-fold), both with potential roles in the nervous system. Most of the remaining downregulated proteins lack established neurological functions or remain uncharacterized in zebrafish. As expected, the zGluN2Aa subunit (A0A8M9PXD6) was absent in *grin2Aa⁻/⁻* samples.

**Figure 8.**
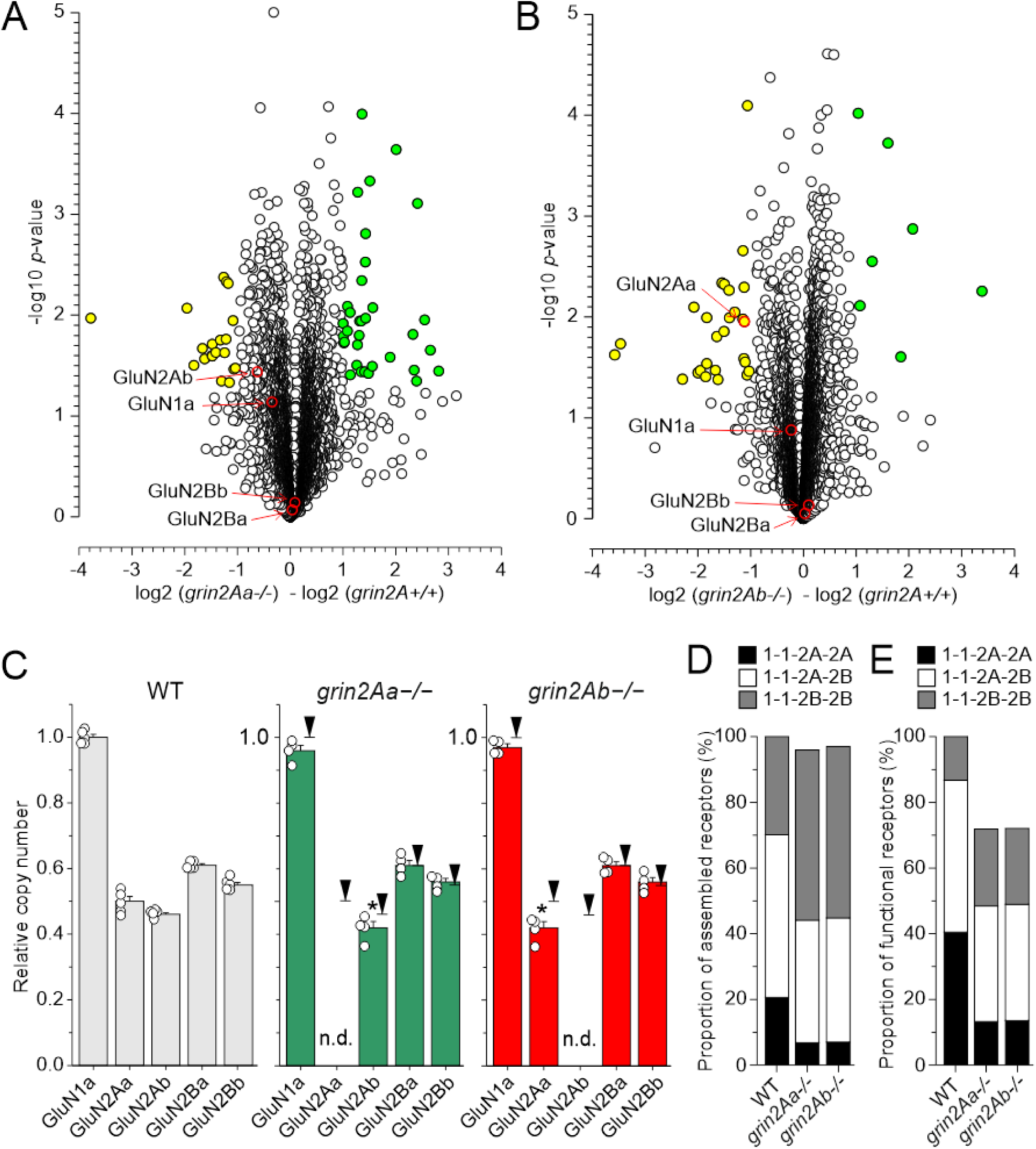
Proteomic analysis of the optic tectum from *grin2A^+/+^*, *grin2Aa^−/−^*, and *grin2Ab^−/−^* adult fish. The volcano plots show proteomics data. The abscissa displays negative (downregulated) and positive (upregulated) fold-changes in the ratio of protein levels found in *grin2Aa*^−/−^ relative to *grin2A^+/+^* (A) and *grin2Ab*^−/−^ relative to *grin2A^+/+^* (B). The statistical significance (-log of *p*-values; indicated on the ordinate) was assessed using two-sample test with the level of significance determined using permutation-based FDR (see Methods for details). Yellow symbols show proteins with significantly downregulated expression and green symbols show proteins with significantly upregulated expression (see Tables S3 and S3 for details of proteins with altered expression). C, The plots show the relative copy abundance number (CAN) of GluN1a, GluN2Aa, GluN2Ab, GluN2Ba, and GluN2Bb expressed in *grin2A^+/+^* (*on the left*), *grin2Aa*^−/−^ (*in the middle*), and *grin2Ab*^−/−^ fish (*on the right*). CAN of individual NMDAR subunits was normalized with respect to CAN of GluN1a assessed in the *grin2A^+/+^* fish. Black arrows indicate relative expression in the *grin2A^+/+^* fish; n.d. stands for not detected. One-way ANOVA was used to assess the significance in the relative CAN of NMDAR subunits. Relative CAN of GluN2Aa and GluN2Ab were significantly changed when compared to *grin2A^+/+^* (*). (D, E) Relative CAN of GluN1a, GluN2Aa, GluN2Ab, GluN2Ba, and GluN2Bb obtained in *grin2A^+/+^*, *grin2Aa*^−/−^ and *grin2Ab*^−/−^ fish were used to calculate (see Supplementary information on calculations) the expected proportions of assembled (D) and functional (E) NMDARs of different subunit compositions, including diheteromeric GluN1a/GluN2A (1-1-2A-2A) and GluN1a/GluN2B receptors (1-1-2B-2B), and triheteromeric GluN1a/GluN2A/GluN2B receptors (1-1-2A-2B).

Bi-allelic deletion of *grin2Ab* significantly altered the expression of 34 proteins compared to *grin2A^+/+^* controls (Fig. 8A, Table S3). The most prominently upregulated protein was C-reactive protein (CRP; Q0IIP8), which showed a 10.5-fold increase. CRP is typically produced in the liver during inflammation (Sproston and Ashworth, 2018); however, no pathogens were identified in microbiome analyses, leaving the cause of this upregulation unclear. The most downregulated proteins (0.08–0.24-fold) included chymotrypsin B1 precursor (Q7SX97) and two uncharacterized proteins (A0A8M2B4C1, A9JRA9), none of which have established links to the glutamatergic system or brain function. As expected, the zGluN2Ab subunit (F1QDE5) was not detected in *grin2Ab⁻/⁻* fish, and the expression level of the zGluN2Aa subunit was also significantly reduced (Fig. 8B, Table S4). NMDAR subunits zGluN1b, zGluN2Ca/b, and zGluN2Da/b were not detected at quantifiable levels in any of the genotypes examined.

The role of NMDA receptors in neuronal excitability, synaptic transmission, and the regulation of neurogenesis, neurite outgrowth, and synapse formation is well established (Hansen et al., 2021). To assess the potential impact of zGluN2A subunit loss on synaptogenesis, we analyzed the abundance of 55 selected presynaptic proteins in the adult zebrafish brain. No significant differences were observed in the expression levels of proteins involved in vesicle exocytosis and vesicular glutamate transport among *grin2A^+/+^*, *grin2Aa^−/−^*, and *grin2Ab^−/−^* fish (p = 0.9973; one-way ANOVA; Table S5). On average, the expression ratio of these presynaptic proteins was 0.9948 for *grin2Aa^−/−^ vs. grin2A^+/+^*, and 1.0052 for *2Ab^−/−^ vs. grin2A^+/+^*, indicating preserved presynaptic protein composition.

Relative copy abundance numbers (CAN) per cell were used to quantify changes in the expression of individual NMDAR subunits in mutant zebrafish. This approach relies on the mass spectrometry signal of histones, which correlates with DNA content and, by extension, cell number (see Methods). In *grin2A^+/+^* fish, the relative CAN of zGluN2Aa, zGluN2Ab, zGluN2Ba, and zGluN2Bb subunits ranged from 0.46 to 0.61, normalized to zGluN1a levels (Fig. 8C). In *grin2Aa^−/−^* fish, zGluN2Aa expression was absent, and zGluN2Ab levels were reduced by 10% compared to *grin2A^+/+^*. The CAN of zGluN1a, zGluN2Ba, and zGluN2Bb remained unchanged. Conversely, in *grin2Ab^−/−^* fish, zGluN2Ab was undetectable, and zGluN2Aa levels were significantly reduced by 16%. No changes were observed in the CAN of zGluN1a, zGluN2Ba, or zGluN2Bb in these mutants.

Quantitative data from MS analysis were used to estimate the abundance and relative proportions of NMDARs containing zGluN2A, zGluN2B, or both subunits in mutant zebrafish. These calculations (see Supplementary Information for details) were based on the following assumptions: NMDARs are tetrameric complexes composed of two zGluN1 subunits and two zGluN2 subunits, which may include zGluN2Aa, zGluN2Ab, zGluN2Ba, or zGluN2Bb (Al-Hallaq et al., 2007; Luo et al., 1997; Tovar et al., 2013; Rauner and Kohr, 2011). It was further assumed that zGluN1 subunits associate with zGluN2 subunits randomly and with equal preference (Kellermayer et al., 2018), although alternative models suggest preferential assembly patterns (Al-Hallaq et al., 2007).

The proportion of diheteromeric zGluN1a/zGluN2A receptors (containing either zGluN2Aa or zGluN2Ab) is predicted to decrease from 20.5% in *grin2A^+/+^* fish to 6.7% and 6.9% in *grin2Aa^−/−^* and *grin2Ab^−/−^* fish, respectively (Fig. 8D). Conversely, the proportion of diheteromeric zGluN1a/zGluN2B receptors is expected to rise from 29.9% in *grin2A^+/+^*to 52.0% and 52.3% in *grin2Aa^−/−^* and *grin2Ab^−/−^*fish. The fraction of triheteromeric zGluN1a/zGluN2A/zGluN2B receptors (containing both zGluN2A and zGluN2B paralogs) is predicted to change only modestly, from 49.6% in *grin2A^+/+^* to 37.3% and 37.9% in *grin2Aa^−/−^* and *grin2Ab^−/−^*fish, respectively (Fig. 8D). Importantly, although the total number of assembled NMDARs is not expected to be reduced—given the excess of zGluN2 subunits relative to zGluN1a (Fig. 8C, D)—this does not necessarily translate to functional receptors. Notably, zGluN2Ba-containing receptors (XM_009299013.2) are non-functional based on electrophysiological data. In *grin2A^+/+^* fish, four GluN2 subunit types are present, with one (zGluN2Ba) forming non-functional receptors. In *grin2Aa^−/−^* and *grin2Ab^−/−^* fish, only three GluN2 subunit types remain, still including the non-functional zGluN2Ba. Thus, the total number of functional NMDARs is predicted to be reduced in *grin2Aa^−/−^* and *grin2Ab^−/−^* fish relative to *grin2A^+/+^* (Fig. 8E). This reduction, combined with shifts in receptor subunit composition (Table 1), may significantly impact synaptic signaling and circuit function in zebrafish lacking functional *grin2A* paralogs.

## DISCUSSION

Using zebrafish as a novel model of NMDAR-related neurodevelopmental pathology, we investigated the molecular and behavioral consequences of disrupting *grin2A* paralogs. We demonstrate that the functional properties of recombinant zebrafish NMDARs are subunit-dependent, following a pattern consistent with that observed in human and rodent receptors. Expression of *grin2A* paralogs begins early in development, and disruption of either *grin2Aa* or *grin2Ab* results in increased locomotor activity in larvae exposed to a novel environment. Proteomic analyses indicate limited zGluN1 availability relative to the abundance of zGluN2 subunits in wild-type fish. In *grin2Aa* or *grin2Ab* mutants, the relative proportion of zGluN2B-containing NMDARs is increased, despite the absence of compensatory *grin2B* upregulation. Together, these findings align with and extend insights from rodent models and contribute to a deeper understanding of GRIN2A loss-of-function mechanisms in neurodevelopmental disorders.

### Development and mRNA

Our findings demonstrate that the expression of *grin2Aa* and *grin2Ab* mRNA in the zebrafish nervous system is both developmentally and anatomically regulated (Fig. 2). Previous studies have shown that *grin1* and *grin2* paralogs exhibit differential expression patterns during early zebrafish development (Cox et al., 2005; Zoodsma et al., 2020; Zoodsma et al., 2022). For example, *grin1a* is expressed in the telencephalon, retina, and spinal cord at 1 dpf, whereas *grin1b* is not detectable at this stage. By 2–5 dpf, both *grin1a* and *grin1b* become broadly expressed throughout the brain and retina (Cox et al., 2005; Zoodsma et al., 2020; Griffin et al., 2021). Similarly, *grin2Aa* and *grin2Ab* mRNA are not detected at 1 dpf but begin to be expressed by 2 dpf, particularly in the retina (Cox et al., 2005), consistent with our results. Their expression increases progressively over the subsequent days and becomes more widespread across brain regions and the retina, though it remains absent from the spinal cord. Expression of *grin2B* paralogs also appears to begin after 1 dpf, with transcripts detected at 2, 3, and 5 dpf in previous reports (Cox et al., 2005; Zoodsma et al., 2022), in agreement with our findings (Fig. S1). Together, these data indicate that the onset of *grin1* and *grin2* gene expression occurs early in zebrafish development and supports the idea that the role of NMDAR signaling in neurodevelopment is evolutionarily conserved.

### Behavior of larvae

Our behavioral analysis shows that deletion of *grin2Aa* and *grin2Ab* enhances locomotor activity in the novel environment primarily by an increase in swim bout frequency. Additionally, we find that the deletion of *grin2Aa vs. grin2Ab* has opposite effects on thigmotaxis (Fig. 4H and 5D-F). The behavioral assay used in our study closely resembles the larval swim behavior monitoring during acclimation described by Zoodsma et al. (Zoodsma et al., 2020; Zoodsma et al., 2022), although our data suggest a longer adaptation period. Notably, (Zoodsma et al., 2020) reported that zebrafish larvae with homozygous deletion of both *grin1a* and *grin1b* retain the ability to perform coordinated swimming and exhibit hyperactivity during initial exposure to a novel swim arena. These findings, together with our own, suggest that NMDARs are not essential for basic locomotor function, but rather contribute to behavioral adaptation in response to environmental novelty.

Our observation of hyperactivity in *grin2Aa* and *grin2Ab* mutant larvae in a novel environment suggests a role for zGluN2A paralogs in behavioral adaptation. In contrast, locomotion in *grin2Bb^−/−^* larvae was indistinguishable from that of control *grin2B⁺/⁺* larvae, consistent with previous findings following deletion of both *grin2B* paralogs (Zoodsma et al., 2022). These results support subunit-specific contributions of NMDAR signaling to zebrafish behavior. Notably, while overall locomotion was elevated across *grin2A* single and double mutants, effects on thigmotaxis were divergent (Fig. 4H and 5 D-F). *grin2Aa^+/−^* and *grin2Aa^−/−^* fish spent more time in the center zone, consistent with an anxiolytic phenotype, whereas *grin2Ab^−/−^*fish spent less time in the center, indicative of increased anxiety. Interestingly, larvae with combined deletion of *grin2Aa* and *grin2Ab* exhibited further increased locomotor activity, but the opposing effects on thigmotaxis were abolished (Fig. 4H and 5E, F). These findings suggest that the hyperactivity observed in *grin2A* mutants is not driven primarily by anxiety, but rather reflects distinct and possibly opposing roles of *grin2Aa* and *grin2Ab* in modulating locomotor adaptation and anxiety-related behavior.

Several factors may contribute to the behavioral differences observed in the knockout fish. At the molecular level, zGluN2Ab shares high homology with rat GluN2A at the distal end of the C-terminal domain (CTD), including the canonical PDZ-binding motif (PSIESDV) that mediates interaction with PSD-95 (Kornau et al., 1995). In contrast, zGluN2Aa contains a variant motif (PSLESDV), more commonly found in rodent GluN2C/D subunits. This single amino acid substitution may influence the synaptic localization and mobility of the zGluN2A paralogs (Bard and Groc, 2011), potentially contributing to their divergent roles in circuit function and behavior. Further research is needed to delineate the specific contributions of NMDAR signaling in distinct zebrafish neural circuits (Wang and McLean, 2014; Berg et al., 2023; Jesuthasan, 2012), and to determine how the disruption of different *grin* genes results in distinct behavioral phenotypes.

### Molecular consequences of GRIN2A loss of function

Approximately 20% of disease-associated *GRIN2A* variants are nonsense or frameshift mutations introducing premature stop codons into the GluN2A subunit sequence, and these are frequently associated with various forms of epilepsy (Santos-Gómez et al., 2020). For nonsense variants located within the ATD, ABD, or TMD, the resulting truncated subunits are unlikely to assemble into functional NMDARs (Korinek et al., 2024). However, the mechanism by which loss-of-function *GRIN2A* variants—reducing the availability of a key NMDAR subunit—lead to circuit hyperexcitability and epilepsy remains unclear. Notably, premature stop codons introduced near the region targeted in this study have been identified in multiple individuals with epilepsy, including the variants A149SfsTer8, W167Ter, W198Ter, F210LfsTer10, Q218Ter, L334Ter, V529WfsTer22, and S538Ter. All individuals were heterozygous and presented with seizures, with or without comorbid developmental delay or intellectual disability (www.grin-database.de). In our study, the mutant zebrafish GluN2A subunits—truncated at 344 and 244 amino acids for zGluN2Aa and zGluN2Ab, respectively— represent ATD-only fragments incapable of forming functional receptors with zGluN1 (Schorge and Colquhoun, 2003). Behaviorally, zebrafish lacking either or both zGluN2A paralogs (*grin2Aa*^−/−^, *grin2Ab*^−/−^, or *grin2A^−/−^*) exhibited increased locomotor activity in a novel environment (Fig. 4 and 5).

To better understand the paradoxical genetic and physiological consequences of *grin2Aa/b* deletion, we investigated potential compensatory mechanisms. We hypothesized that loss of *grin2Aa* and/or *grin2Ab* leads to reduced levels of zGluN2A subunits, which may be offset by upregulation of other NMDAR subunits—particularly zGluN2B—resulting in altered receptor subunit composition. NMDARs containing GluN2B subunits exhibit slower deactivation kinetics than those containing GluN2A (Paoletti et al., 2013; Wyllie et al., 2013) (Fig. 1B). Evidence from rodent studies suggests that substituting GluN2A with GluN2B can prolong NMDAR activation, indicating that protein-truncating variants in *GRIN2A* may produce a net gain-of-function effect by altering receptor kinetics and potentially enhancing excitatory signaling (Camp et al., 2023).

Our RT-qPCR analysis shows that *grin2Ab* mRNA is significantly downregulated in *grin2Ab* mutant fish (Fig. 7B), consistent with nonsense-mediated mRNA decay (NMD)—a surveillance mechanism that promotes degradation of transcripts containing premature stop codons (Nagy and Maquat, 1998; Chang et al., 2007; Rebbapragada and Lykke-Andersen, 2009). Similar downregulation of mutant *grin1a*, *grin2Ba*, and *grin2Bb* transcripts has been reported in corresponding zebrafish models (Zoodsma et al., 2020; Zoodsma et al., 2022). Despite the involvement of NMD, we did not observe robust compensatory upregulation of other NMDAR subunits, particularly the *grin2B* paralogs (Fig. 7B), which contrasts with the increased expression of *grin1a*, *grin1b*, and *grin2Ab* observed in *grin2B^−/−^* larvae (Zoodsma et al., 2022). It is important to note, however, that our analysis assessed mRNA expression at the whole-tissue level and may not capture cell-type-specific changes in NMDAR subunit expression or function within neural circuits involved in locomotor behavior and thigmotaxis.

Proteomic analysis revealed that zGluN2Aa/b and zGluN2Bb/a paralogs are expressed at comparable levels in the optic tectum of adult *grin2A^+/+^* zebrafish, with each GluN2 subunit reaching approximately 50% of the expression level of the zGluN1a subunit (Fig. 8D). Notably, we did not detect zGluN1b protein, despite the confirmed presence of *grin1b* mRNA in both larval and adult stages (Figs. S1 and S3). Although previous studies suggest that *grin1b* deletion produces a milder phenotype than *grin1a* deletion, they nonetheless support a functional role for the zGluN1b subunit (Zoodsma et al., 2020). Our modeling assumes negligible zGluN1b contribution to NMDAR assembly. Under this assumption, the majority of available zGluN1a subunits are predicted to be incorporated into functional receptors, while a substantial fraction of zGluN2 subunits remains unassembled due to limiting zGluN1 availability. This imbalance suggests that in zebrafish harboring nonsense mutations in *grin2Aa* or *grin2Ab*, the proportion of zGluN2B-containing receptors may increase without requiring compensatory *grin2B* transcriptional upregulation or increased zGluN2B protein abundance. Given the slower deactivation kinetics of zGluN2B-containing NMDARs (Fig. 1B), this shift in receptor subunit composition may enhance synaptic excitation and contribute to altered circuit function in mutant fish.

This study provides new insights into the effects of *grin2A* nonsense mutations in zebrafish. Our findings underscore the functional similarities between zebrafish and mammalian NMDARs and extend observations from rodent models of *GRIN2A* loss of function. These results support the use of zebrafish as a valuable model for investigating the mechanisms underlying *GRIN*-related neurodevelopmental disorders, an essential step toward developing effective therapies.

## Abbreviations

ABD: agonist-binding domain
ANOVA: analysis of varinace
ATD: amino-terminal domain
CAN: copy abundance number
CTD: C-terminal domain
dpf: days post fertilization
ECS: extracellular solution
FDR: false discovery rate
GFP: green fluorescent protein
HEK293T: human embryonic kidney 293T
hGluN: human variant of GluN subunit
ISH: *in-situ* hybridization
LSD: least significant difference
M1 – M4: membrane helices 1 -4
MBD: mean bout distance
MBF: mean bout frequency
MTD: mean travelled distance
NMDAR: N-methyl-D-aspartate receptor
RT-qPCR: reverse transcription PCR
SEM: standard error of the mean
TMD: transmembrane domain
τw: weighted tau of double-exponential fit
zGluN: zebrafish variant of GluN subunit

## Acknowledgements

This work was supported by the Czech Science Foundation (GACR): 23-04922S, GAUK project No. 410122, and Research Project of the CAS RVO: 67985823. We thank Prof. Reinhard Köster and Dr. Franz Vauti (Technical University Braunschweig, Germany) for advice on the whole mount *in-situ* hybridization technique, Dr. Björn Schuster and Mgr. Ivana Dobiasovska (Institute of Molecular Genetics CAS, Prague) for help with the preparation of *grin* zebrafish lines, and staff of the fish facility (Institute of Molecular genetics CAS, Prague) and Mgr. Romana Markova (Institute of Physiology CAS, Prague) for excellent technical assistance.

## Author contributions

Vera Abramova prepared mutant fish strains, did *in-situ* hybridization and behavioral studies. Bohdan Kysilov, Mark Dobrovolski, Barbora Hrcka Krausova, Klevinda Fili and Fatma Elzahraa S. Abdel Rahman performed electrophysiological experiments and data analysis. Vera Abramova, Paulina Bozikova, Jiri Cerny, Tereza Smejkalova, Miloslav Korinek, Ales Balik and Ladislav Vyklicky performed data analysis. Eni Tomovic and Ales Balik performed RT-qPCR. Tereza Smejkalova, Miloslav Korinek and Ladislav Vyklicky wrote the manuscript. Ladislav Vyklicky supervised the project, designed the research and conceptualized the data. All authors discussed the results, commented on the manuscript and approved it.

## Data availability

Data is available from the corresponding author upon reasonable request.

## Ethics approval

Zebrafish (*Danio rerio*) were handled and maintained in accordance with Directive 2010/63/EU on the protection of animals used for scientific purposes, and national, and institutional guidelines.

## SUPPLEMENTARY INFORMATION

### Supplementary Tables

**Table S1.**
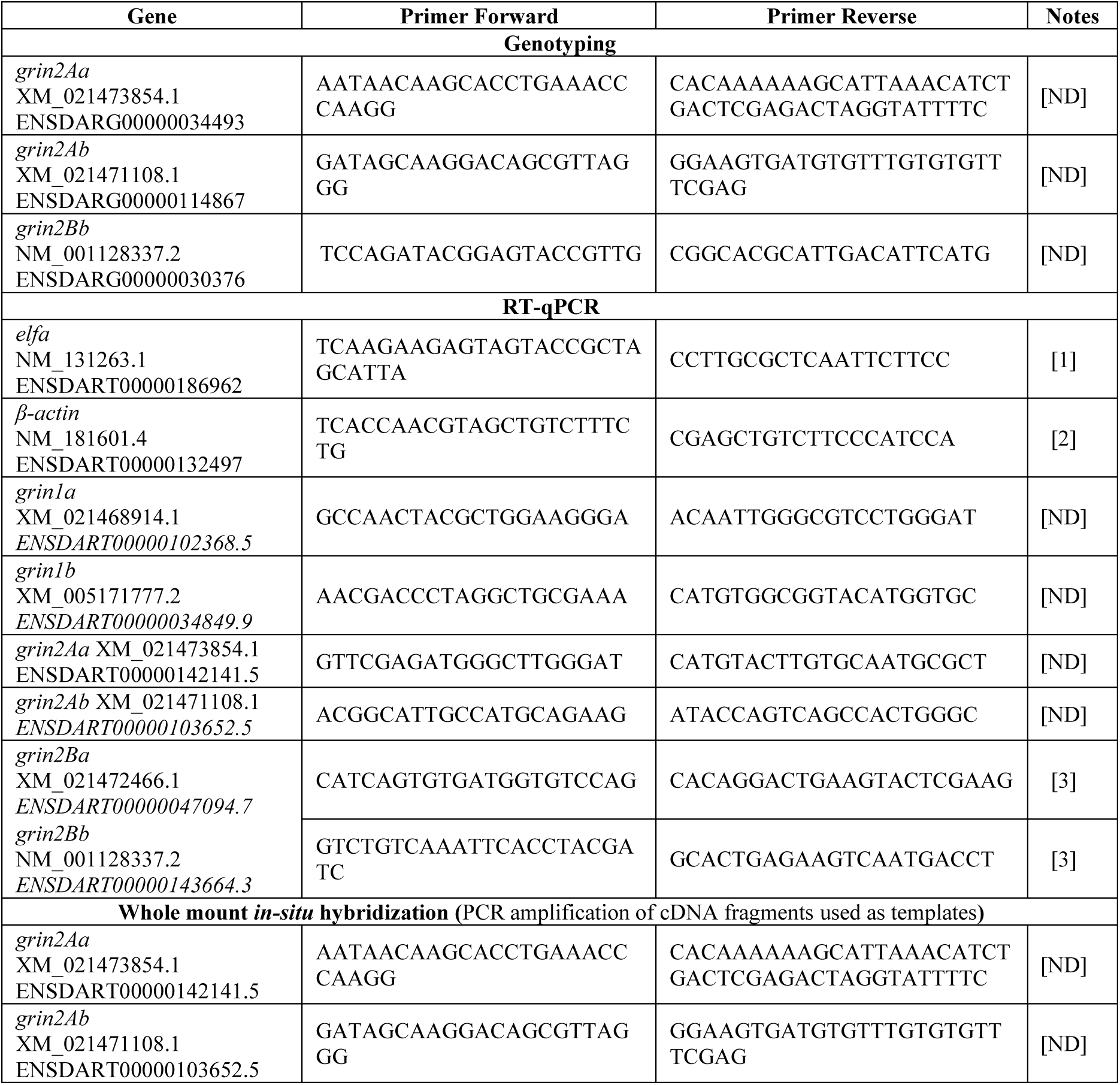
List of primers used for zebrafish genotyping, RT-qPCR and PCR amplification of cDNA fragments used as templates for *in-situ* hybridization probe synthesis. [ND], newly designed; [1], (Moussavi Nik et al., 2014); [2], (Lang et al., 2016); [3], (Zoodsma et al., 2022).

**Table S2.**
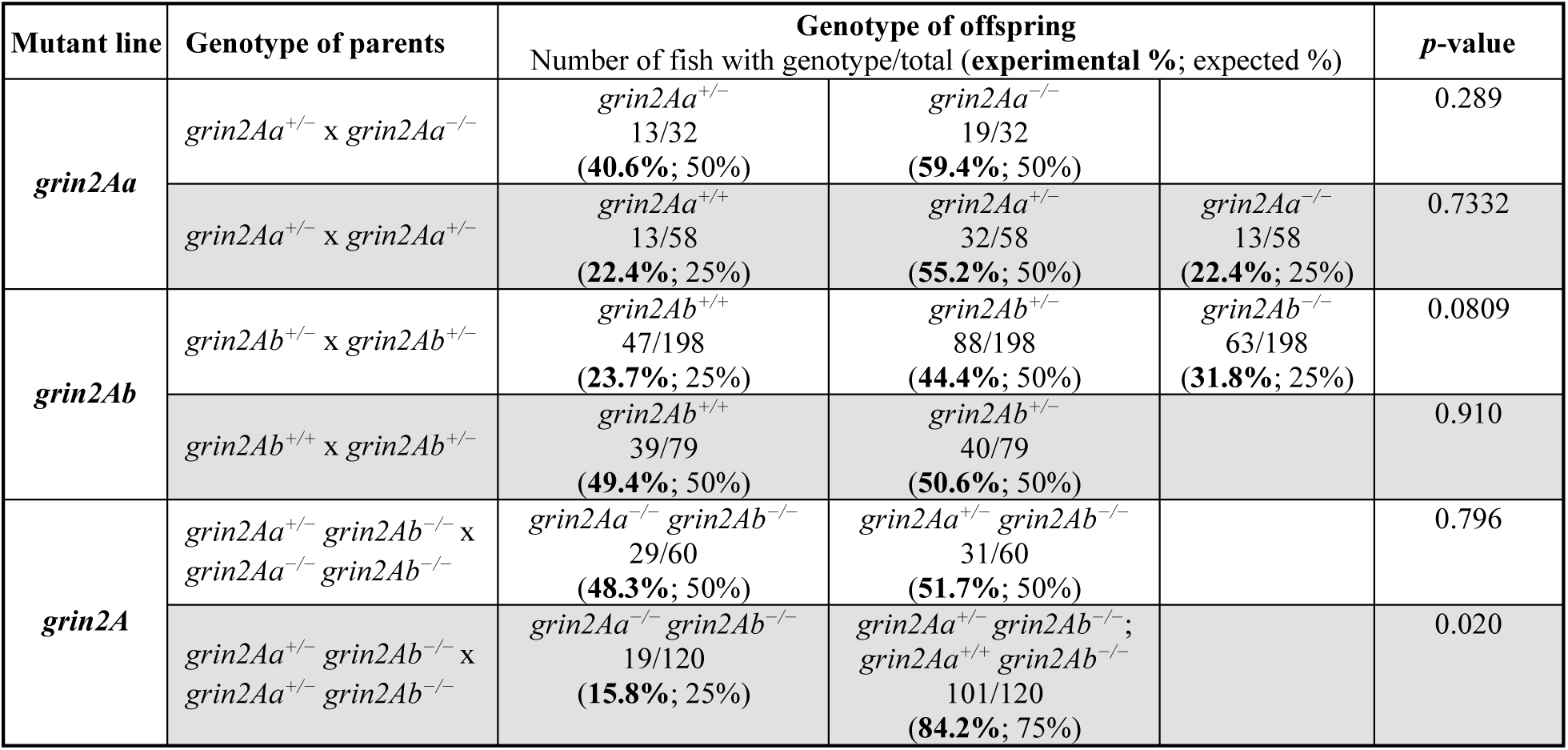
Viability of larvae with mono- and bi-allelic deletion of *grin2Aa, grin2Ab,* or both genes at 6 dpf. Larvae are from different intercrosses between *grin2Aa*, *grin2Ab*, and double mutants. *χ^2^* was used to assess the significance.

**Table S3.**
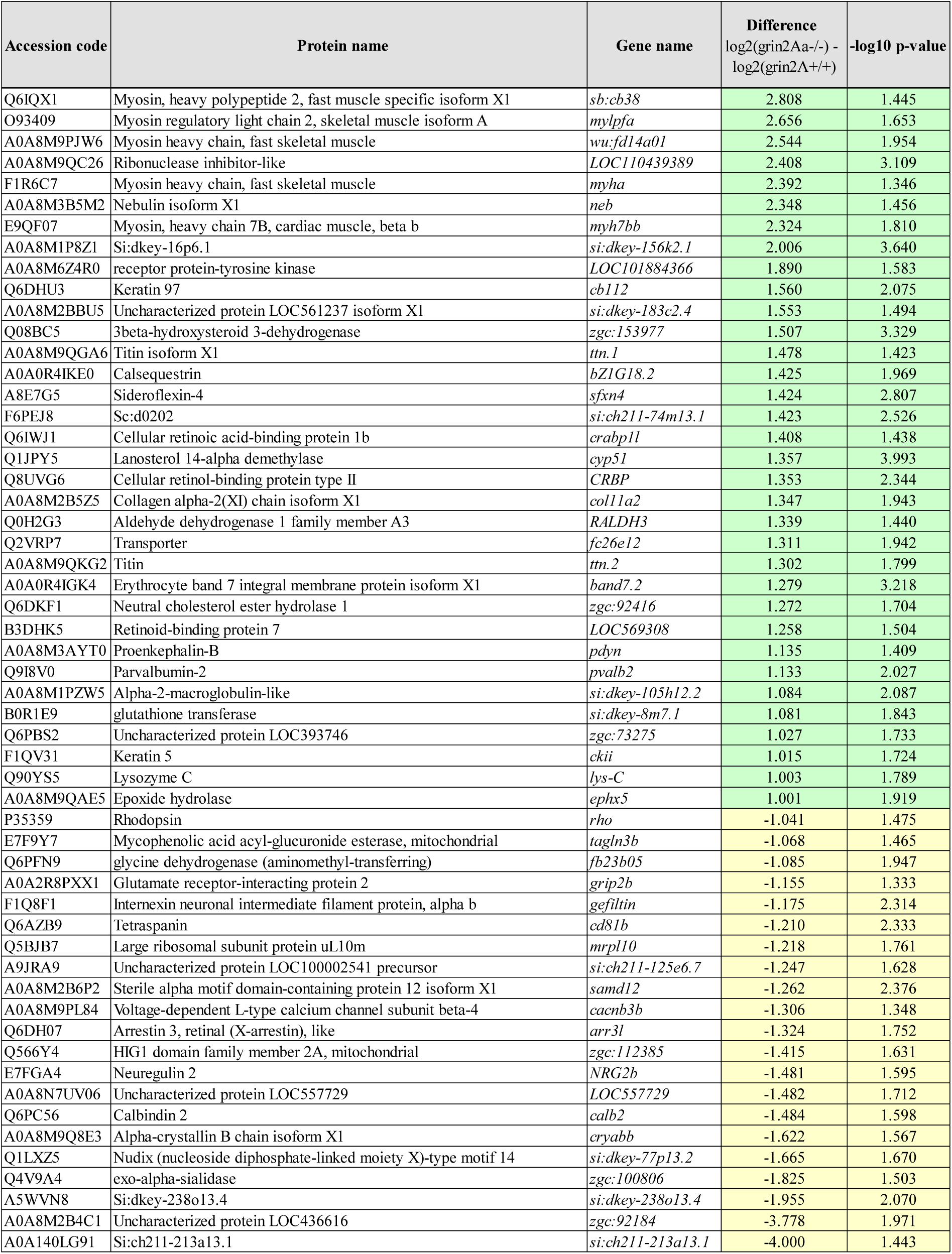
Proteomic analysis of brain tissue from adult *grin2Aa^-/-^* zebrafish. Proteins showing increased (green) or decreased (yellow) expression levels compared to *grin2A^+/+^* zebrafish are listed.

**Table S4.**
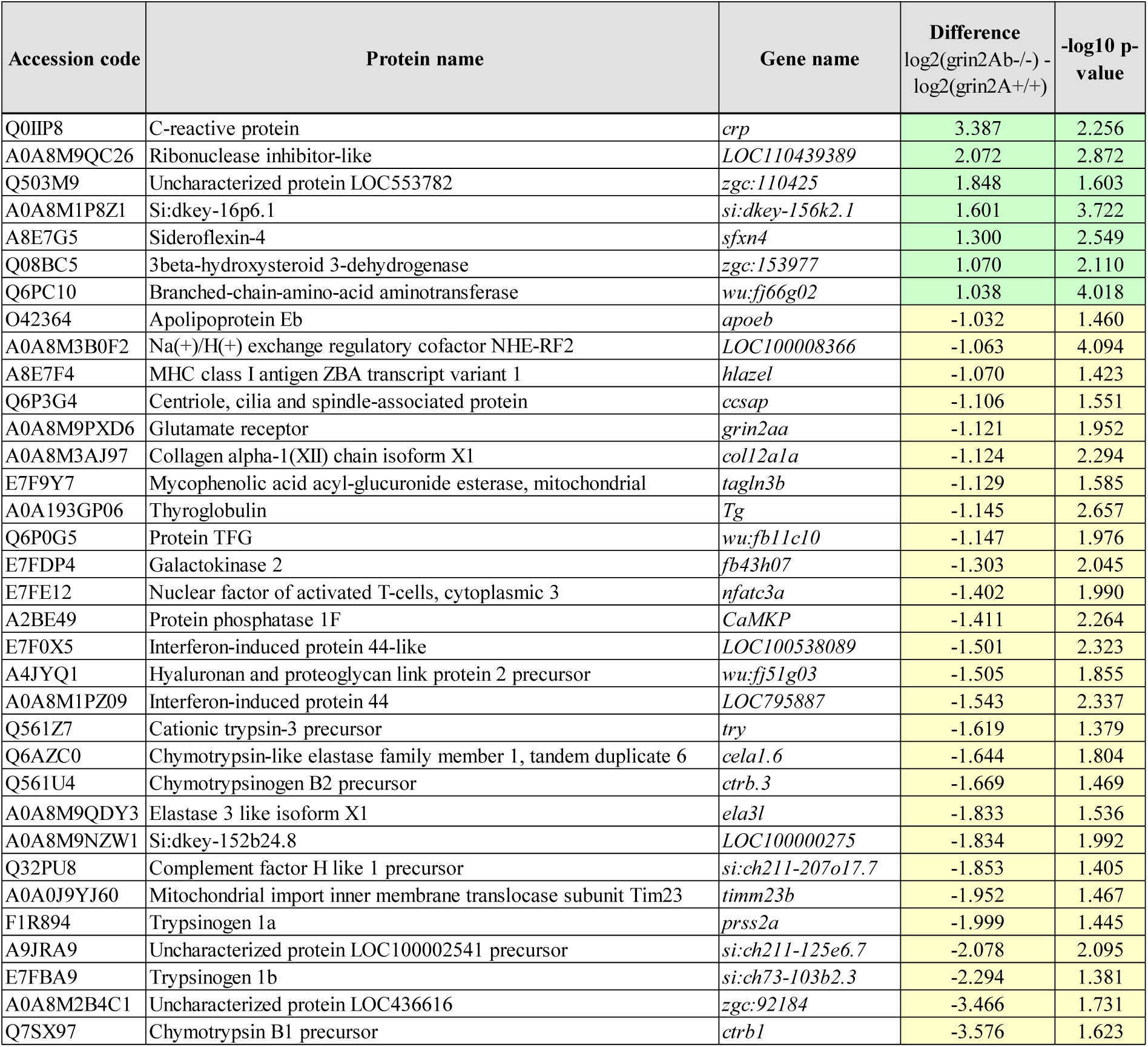
Proteomic analysis of brain tissue from adult *grin2Ab^-/-^* zebrafish. Proteins showing increased (green) or decreased (yellow) expression levels compared to *grin2A^+/+^* zebrafish are listed.

**Table S5.**
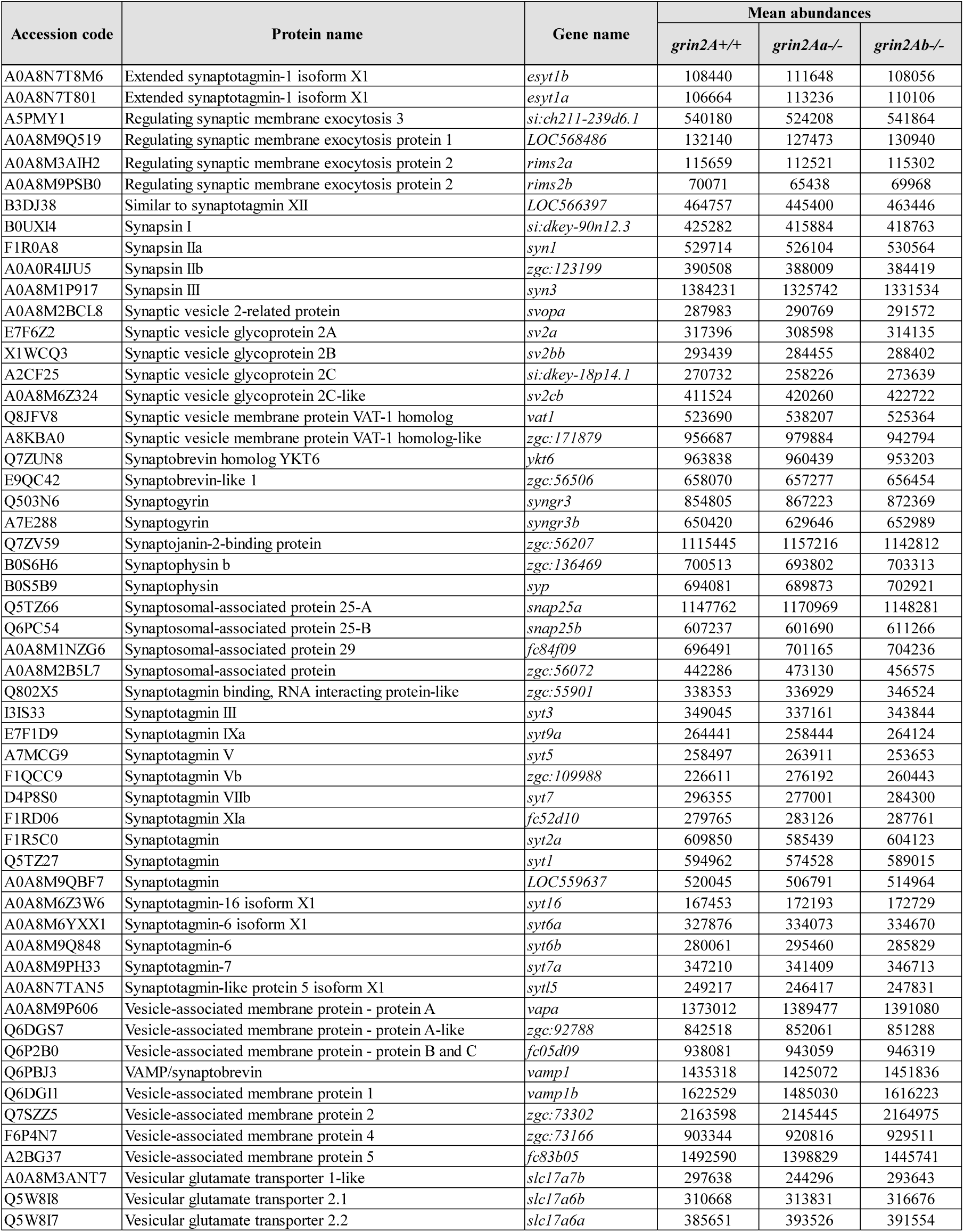
Proteomic analysis of selected presynaptic proteins. Mean abundances of selected presynaptic proteins found in the brain of *grin2A^+/+^* (*n* = 5), *grin2Aa*^−/−^ (*n* = 4), and *grin2Ab*^−/−^ (*n* = 4) adult zebrafish. One-way ANOVA showed no genotype differences in the abundance levels (*p* = 0.9973).

**Figure S1.**
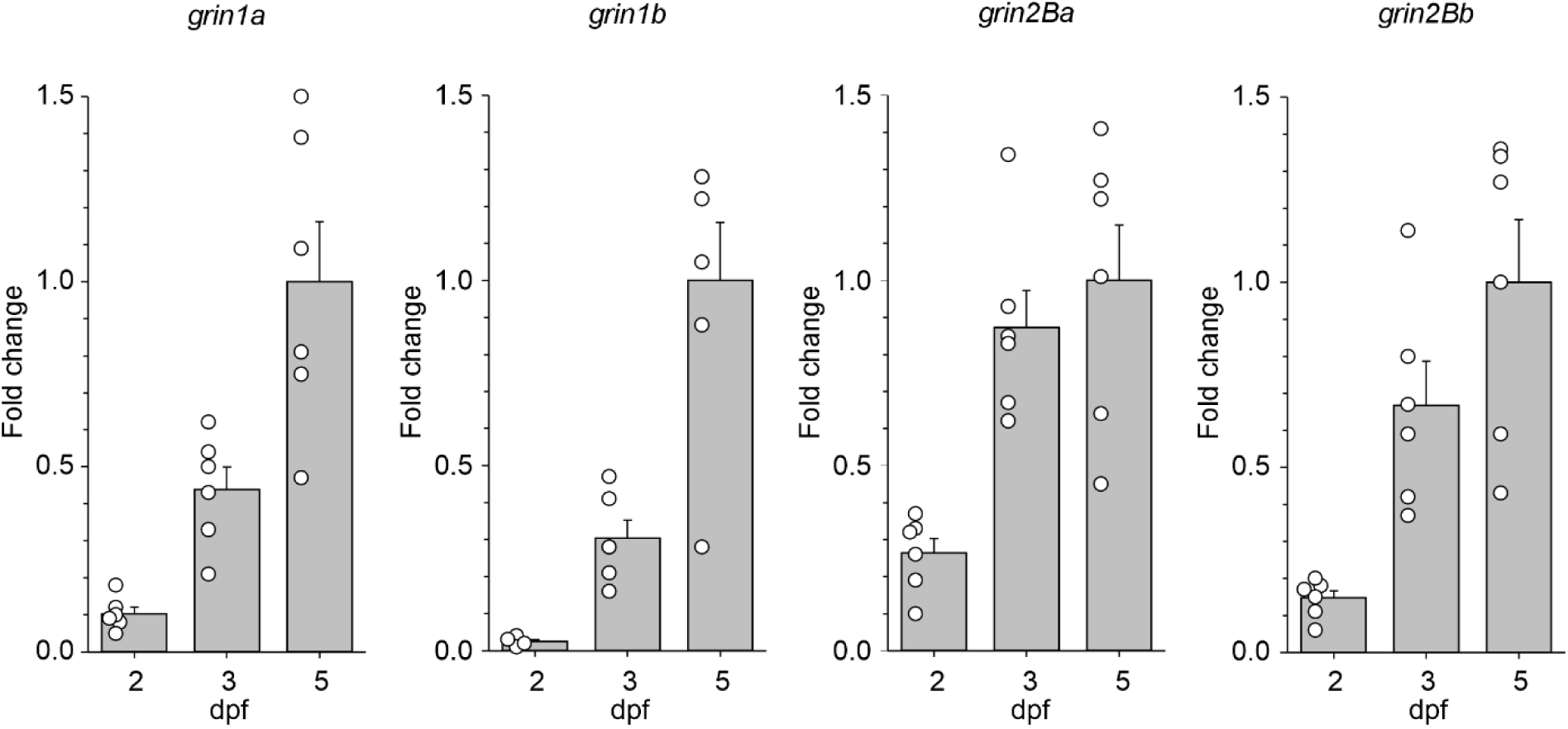
The mRNA expression levels of *grin1* and *grin2B* paralogs in the zebrafish nervous system. The expression levels of *grin1a*, *grin1b*, *grin2Ba*, and *grin2Bb* mRNA in heads of zebrafish larvae at 2, 3, and 5 dpf were analyzed by RT-qPCR. Data were normalized to the expression of the corresponding gene at 5 dpf; *grin1a*: *n* = 6, 6, 6; *grin1b*: *n* = 6, 5, 5; *grin2Ba*: *n* = 6, 6, 6; *grin2Bb*: *n* = 6, 6, 6.

**Figure S2.**
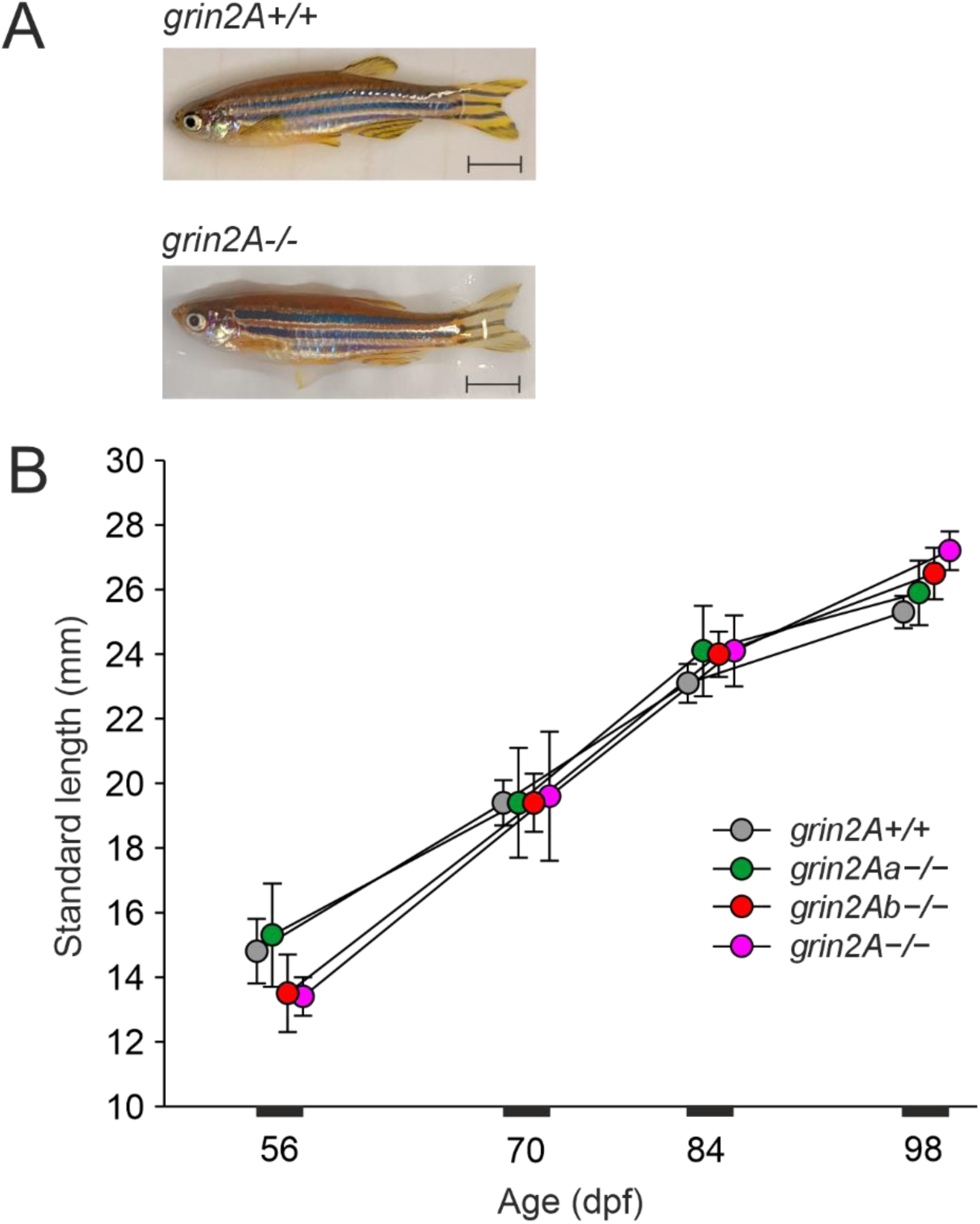
During development, *grin2Aa*^−/−^, *grin2Ab*^−/−^, *grin2A*^−/−^ mutant fish show normal growth. A, Representative photographs of *grin2A^+/+^* and *grin2A*^−/−^ fish at 98 dpf. Scale bar, 5 mm. B, Standard length (from snout to the tail peduncle) comparison of *grin2A^+/+^*(*n* = 12), *grin2Aa*^−/−^ (*n* = 6), *grin2Ab*^−/−^ (*n* = 9), and *grin2A*^−/−^ (*n* = 4). One-way ANOVA showed no differences in the standard length among the genotypes at 56 (*p* = 0.654), 70 (*p* = 0.999), 84 (*p* = 0.784), and 98 (*p* = 0.395) dpf.

**Figure S3.**
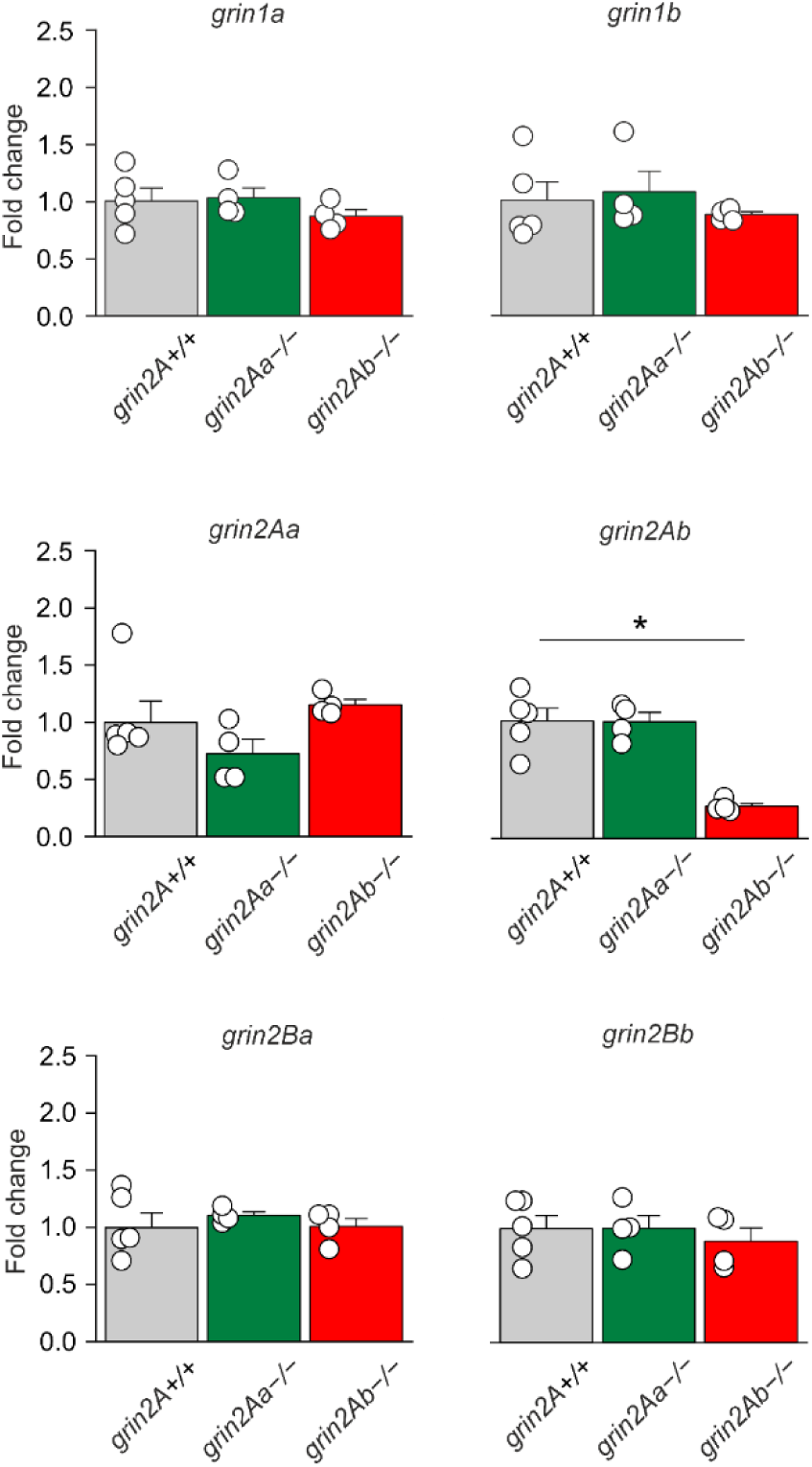
Effects of *grin2Aa* or *grin2Ab* deletion on *grin* mRNA expression in the adult zebrafish. The relative RT-qPCR expression levels of *grin1a, grin1b, grin2Aa, grin2Ab, grin2Ba* and *grin2Bb* mRNA assessed in optic tectum of *grin2A^+/+^*, *grin2Aa^−/−^*, and *grin2Ab^−/−^*fish. One-way ANOVA was used to assess differences in the relative expression of each gene, followed by multiple comparisons *versus* corresponding gene in the *grin2A^+/+^*fish (Dunnett’s method), with significance indicated by *. Sample sizes for *grin2A^+/+^*, *grin2Aa^−/−^*, and *grin2Ab^−/−^*, respectively, and ANOVA results are as follows: *grin1a*: *n* = 5, 4, 4, *p* = 0.422; *grin1b*: *n* = 5, 4, 4, *p* = 0.477; *grin2Aa*: *n* = 5, 4, 4, *p* = 0.04; *grin2Ab*: *n* = 5, 4, 4, *p* < 0.001; *grin2Ba*: *n* = 5, 4, 4, *p* = 0.760; and *grin2Bb*: *n* = 5, 4, 4, *p* = 0.747.

**Figure S4.**
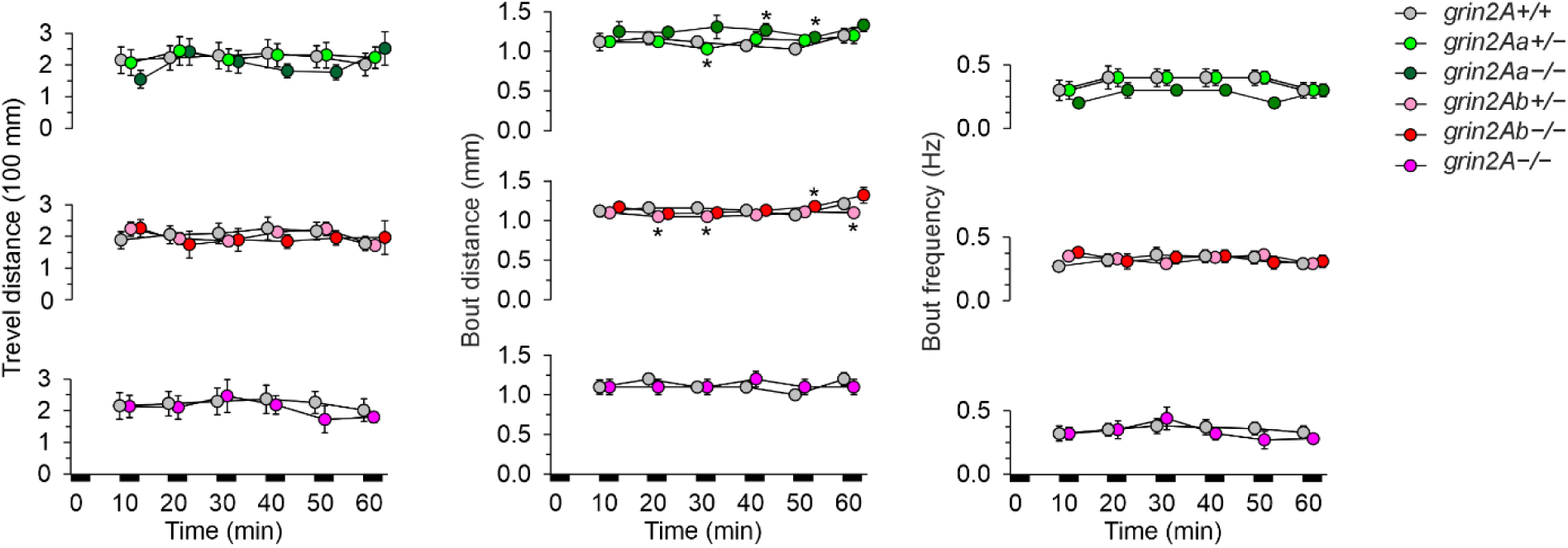
Steady-state swimming parameters. assessed 2–3 h after the larvae were placed in the experimental chambers. * indicates significant differences in genotype as assessed by ANOVA followed by LSD *post-hoc* test (*grin2A^+/+^*: *n* = 258; *grin2Aa^+/−^*: *n* = 204; *grin2Aa^−/−^*: *n* = 95; *grin2Ab^+/−^*: *n* = 428; *grin2Ab^−/−^*: *n* = 119; *grin2A^−/−^*: *n* = 77). *grin2Aa^+/−^* and *grin2Aa^−/−^ versus grin2A^+/+^*: locomotor activity: *p* = 0.261; bout frequency: *p* = 0.143; bout distance: *p* < 0.001; *grin2Ab^+/−^* and *grin2Ab^−/−^ versus grin2A^+/+^*: locomotor activity: *p* = 0.954; bout frequency: *p* = 0.84; bout distance: *p* < 0.001; *grin2A^−/−^ versus grin2A^+/+^*: locomotor activity: *p* = 0.253; bout frequency: *p* = 0.173; bout distance: *p* = 0.8. No time differences in the locomotor activity, bout frequency, and bout distance of *grin2A^+/+^*, *grin2Aa^+/−^*, *grin2Aa^−/−^*, *grin2Ab^+/−^*, *grin2Ab^−/−^*, and *grin2A^−/−^* were found using ANOVA.

