## Supplementary Tables and Figures for "Disruption of *grin2A*, an epilepsy-associated gene, produces altered spontaneous swim behavior in zebrafish"

**Table of Contents:**

**Table S1.** List of primers used for zebrafish genotyping, RT-qPCR and *in-situ* hybridization probe cloning
**Table S2.** Viability of larvae with mono- and bi-allelic deletion of *grin2Aa*, *grin2Ab*, or both genes at 6 dpf
**Table S3.** Proteomic analysis of brain tissue from adult *grin2Aa*$^{-/-}$ zebrafish.
**Table S4.** Proteomic analysis of brain tissue from adult *grin2Ab*$^{-/-}$ zebrafish
**Table S5.** Proteomic analysis of selected presynaptic proteins
**Figure S1.** The mRNA expression levels of *grin1* and *grin2B* paralogs in the zebrafish nervous system
**Figure S2.** During development, *grin2Aa*$^{-/-}$, *grin2Ab*$^{-/-}$, *grin2A*$^{-/-}$ mutant fish show normal growth
**Figure S3.** Effects of *grin2Aa* or *grin2Ab* deletion on *grin* mRNA expression in the adult zebrafish
**Figure S4.** Steady-state swimming parameters

### Supplementary Tables:

| Gene | Primer Forward | Primer Reverse | Notes |
|---|---|---|---|
| | **Genotyping** | | |
| *grin2Aa*<br>XM_021473854.1<br>ENSDARG00000034493 | AATAACAAGCACCTGAAACC CAAGG | CACAAAAAAGCATTAAACATCT GACTCGAGACTAGGTATTTTC | [ND] |
| *grin2Ab*<br>XM_021471108.1<br>ENSDARG00000114867 | GATAGCAAGGACAGCGTTAG GG | GGAAGTGATGTGTTTGTGTGTT TCGAG | [ND] |
| *grin2Bb*<br>NM_001128337.2<br>ENSDARG00000030376 | TCCAGATACGGAGTACCGTTG | CGGCACGCATTGACATTCATG | [ND] |
| | **RT-qPCR** | | |
| *elfa*<br>NM_131263.1<br>ENSDART00000186962 | TCAAGAAGAGTAGTACCGCTA GCATTA | CCTTGCGCTCAATTCTTCC | [1] |
| *β-actin*<br>NM_181601.4<br>ENSDART00000132497 | TCACCAACGTAGCTGTCTTTC TG | CGAGCTGTCTTCCCATCCA | [2] |
| *grin1a*<br>XM_021468914.1<br>*ENSDART00000102368.5* | GCCAACTACGCTGGAAGGGA | ACAATTGGGCGTCCTGGGAT | [ND] |
| *grin1b*<br>XM_005171777.2<br>*ENSDART00000034849.9* | AACGACCCTAGGCTGCGAAA | CATGTGGCGGTACATGGTGC | [ND] |
| *grin2Aa* XM_021473854.1<br>ENSDART00000142141.5 | GTTCGAGATGGGCTTGGGAT | CATGTACTTGTGCAATGCGCT | [ND] |
| *grin2Ab* XM_021471108.1<br>*ENSDART00000103652.5* | ACGGCATTGCCATGCAGAAG | ATACCAGTCAGCCACTGGGC | [ND] |
| *grin2Ba*<br>XM_021472466.1<br>*ENSDART00000047094.7* | CATCAGTGTGATGGTGTCCAG | CACAGGACTGAAGTACTCGAAG | [3] |
| *grin2Bb*<br>NM_001128337.2<br>*ENSDART00000143664.3* | GTCTGTCAAATTCACCTACGA TC | GCACTGAGAAGTCAATGACCT | [3] |
| | **Whole mount *in-situ* hybridization (**PCR amplification of cDNA fragments used as templates**)** | | |
| *grin2Aa*<br>XM_021473854.1<br>ENSDART00000142141.5 | AATAACAAGCACCTGAAACC CAAGG | CACAAAAAAGCATTAAACATCT GACTCGAGACTAGGTATTTTC | [ND] |
| *grin2Ab*<br>XM_021471108.1<br>ENSDART00000103652.5 | GATAGCAAGGACAGCGTTAG GG | GGAAGTGATGTGTTTGTGTGTT TCGAG | [ND] |

**Table S1 | List of primers used for zebrafish genotyping, RT-qPCR and PCR amplification of cDNA fragments used as templates for *in-situ* hybridization probe synthesis.** [ND], newly designed; [1], (Moussavi Nik et al., 2014); [2], (Lang et al., 2016); [3], (Zoodsma et al., 2022).

| Mutant line | Genotype of parents | **Genotype of offspring** Number of fish with genotype/total (**experimental %**; expected %) | | | *p*-value |
|---|---|---|---|---|---|
| ***grin2Aa*** | *grin2Aa*$^{+/-}$ x *grin2Aa*$^{-/-}$ | *grin2Aa*$^{+/-}$ 13/32 (**40.6%**; 50%) | *grin2Aa*$^{-/-}$ 19/32 (**59.4%**; 50%) | | 0.289 |
| | *grin2Aa*$^{+/-}$ x *grin2Aa*$^{+/-}$ | *grin2Aa*$^{+/+}$ 13/58 (**22.4%**; 25%) | *grin2Aa*$^{+/-}$ 32/58 (**55.2%**; 50%) | *grin2Aa*$^{-/-}$ 13/58 (**22.4%**; 25%) | 0.7332 |
| ***grin2Ab*** | *grin2Ab*$^{+/-}$ x *grin2Ab*$^{+/-}$ | *grin2Ab*$^{+/+}$ 47/198 (**23.7%**; 25%) | *grin2Ab*$^{+/-}$ 88/198 (**44.4%**; 50%) | *grin2Ab*$^{-/-}$ 63/198 (**31.8%**; 25%) | 0.0809 |
| | *grin2Ab*$^{+/+}$ x *grin2Ab*$^{+/-}$ | *grin2Ab*$^{+/+}$ 39/79 (**49.4%**; 50%) | *grin2Ab*$^{+/-}$ 40/79 (**50.6%**; 50%) | | 0.910 |
| ***grin2A*** | *grin2Aa*$^{+/-}$ *grin2Ab*$^{-/-}$ x *grin2Aa*$^{-/-}$ *grin2Ab*$^{-/-}$ | *grin2Aa*$^{-/-}$ *grin2Ab*$^{-/-}$ 29/60 (**48.3%**; 50%) | *grin2Aa*$^{+/-}$ *grin2Ab*$^{-/-}$ 31/60 (**51.7%**; 50%) | | 0.796 |
| | *grin2Aa*$^{+/-}$ *grin2Ab*$^{-/-}$ x *grin2Aa*$^{+/-}$ *grin2Ab*$^{-/-}$ | *grin2Aa*$^{-/-}$ *grin2Ab*$^{-/-}$ 19/120 (**15.8%**; 25%) | *grin2Aa*$^{+/-}$ *grin2Ab*$^{-/-}$; *grin2Aa*$^{+/+}$ *grin2Ab*$^{-/-}$ 101/120 (**84.2%**; 75%) | | 0.020 |

**Table S2 | Viability of larvae with mono- and bi-allelic deletion of *grin2Aa, grin2Ab,* or both genes at 6 dpf.** Larvae are from different intercrosses between *grin2Aa*, *grin2Ab*, and double mutants. $\chi^2$ was used to assess the significance.

| Accession code | Protein name | Gene name | Difference log2(grin2Aa-/-) - log2(grin2A+/+) | -log10 p-value |
|---|---|---|---|---|
| Q6IQX1 | Myosin, heavy polypeptide 2, fast muscle specific isoform X1 | *sb:cb38* | 2.808 | 1.445 |
| O93409 | Myosin regulatory light chain 2, skeletal muscle isoform A | *mylpfa* | 2.656 | 1.653 |
| A0A8M9PJW6 | Myosin heavy chain, fast skeletal muscle | *wu:fd14a01* | 2.544 | 1.954 |
| A0A8M9QC26 | Ribonuclease inhibitor-like | *LOC110439389* | 2.408 | 3.109 |
| F1R6C7 | Myosin heavy chain, fast skeletal muscle | *myha* | 2.392 | 1.346 |
| A0A8M3B5M2 | Nebulin isoform X1 | *neb* | 2.348 | 1.456 |
| E9QF07 | Myosin, heavy chain 7B, cardiac muscle, beta b | *myh7bb* | 2.324 | 1.810 |
| A0A8M1P8Z1 | Si:dkey-16p6.1 | *si:dkey-156k2.1* | 2.006 | 3.640 |
| A0A8M6Z4R0 | receptor protein-tyrosine kinase | *LOC101884366* | 1.890 | 1.583 |
| Q6DHU3 | Keratin 97 | *cb112* | 1.560 | 2.075 |
| A0A8M2BBU5 | Uncharacterized protein LOC561237 isoform X1 | *si:dkey-183c2.4* | 1.553 | 1.494 |
| Q08BC5 | 3beta-hydroxysteroid 3-dehydrogenase | *zgc:153977* | 1.507 | 3.329 |
| A0A8M9QGA6 | Titin isoform X1 | *ttn.1* | 1.478 | 1.423 |
| A0A0R4IKE0 | Calsequestrin | *bZ1G18.2* | 1.425 | 1.969 |
| A8E7G5 | Sideroflexin-4 | *sfxn4* | 1.424 | 2.807 |
| F6PEJ8 | Sc:d0202 | *si:ch211-74m13.1* | 1.423 | 2.526 |
| Q6IWJ1 | Cellular retinoic acid-binding protein 1b | *crabp1l* | 1.408 | 1.438 |
| Q1JPY5 | Lanosterol 14-alpha demethylase | *cyp51* | 1.357 | 3.993 |
| Q8UVG6 | Cellular retinol-binding protein type II | *CRBP* | 1.353 | 2.344 |
| A0A8M2B5Z5 | Collagen alpha-2(XI) chain isoform X1 | *col11a2* | 1.347 | 1.943 |
| Q0H2G3 | Aldehyde dehydrogenase 1 family member A3 | *RALDH3* | 1.339 | 1.440 |
| Q2VRP7 | Transporter | *fc26e12* | 1.311 | 1.942 |
| A0A8M9QKG2 | Titin | *ttn.2* | 1.302 | 1.799 |
| A0A0R4IGK4 | Erythrocyte band 7 integral membrane protein isoform X1 | *band7.2* | 1.279 | 3.218 |
| Q6DKF1 | Neutral cholesterol ester hydrolase 1 | *zgc:92416* | 1.272 | 1.704 |
| B3DHK5 | Retinoid-binding protein 7 | *LOC569308* | 1.258 | 1.504 |
| A0A8M3AYT0 | Proenkephalin-B | *pdyn* | 1.135 | 1.409 |
| Q9I8V0 | Parvalbumin-2 | *pvalb2* | 1.133 | 2.027 |
| A0A8M1PZW5 | Alpha-2-macroglobulin-like | *si:dkey-105h12.2* | 1.084 | 2.087 |
| B0R1E9 | glutathione transferase | *si:dkey-8m7.1* | 1.081 | 1.843 |
| Q6PBS2 | Uncharacterized protein LOC393746 | *zgc:73275* | 1.027 | 1.733 |
| F1QV31 | Keratin 5 | *ckii* | 1.015 | 1.724 |
| Q90YS5 | Lysozyme C | *lys-C* | 1.003 | 1.789 |
| A0A8M9QAE5 | Epoxide hydrolase | *ephx5* | 1.001 | 1.919 |
| P35359 | Rhodopsin | *rho* | -1.041 | 1.475 |
| E7F9Y7 | Mycophenolic acid acyl-glucuronide esterase, mitochondrial | *tagln3b* | -1.068 | 1.465 |
| Q6PFN9 | glycine dehydrogenase (aminomethyl-transferring) | *fb23b05* | -1.085 | 1.947 |
| A0A2R8PXX1 | Glutamate receptor-interacting protein 2 | *grip2b* | -1.155 | 1.333 |
| F1Q8F1 | Internexin neuronal intermediate filament protein, alpha b | *gefiltin* | -1.175 | 2.314 |
| Q6AZB9 | Tetraspanin | *cd81b* | -1.210 | 2.333 |
| Q5BJB7 | Large ribosomal subunit protein uL10m | *mrpl10* | -1.218 | 1.761 |
| A9JRA9 | Uncharacterized protein LOC100002541 precursor | *si:ch211-125e6.7* | -1.247 | 1.628 |
| A0A8M2B6P2 | Sterile alpha motif domain-containing protein 12 isoform X1 | *samd12* | -1.262 | 2.376 |
| A0A8M9PL84 | Voltage-dependent L-type calcium channel subunit beta-4 | *cacnb3b* | -1.306 | 1.348 |
| Q6DH07 | Arrestin 3, retinal (X-arrestin), like | *arr3l* | -1.324 | 1.752 |
| Q566Y4 | HIG1 domain family member 2A, mitochondrial | *zgc:112385* | -1.415 | 1.631 |
| E7FGA4 | Neuregulin 2 | *NRG2b* | -1.481 | 1.595 |
| A0A8N7UV06 | Uncharacterized protein LOC557729 | *LOC557729* | -1.482 | 1.712 |
| Q6PC56 | Calbindin 2 | *calb2* | -1.484 | 1.598 |
| A0A8M9Q8E3 | Alpha-crystallin B chain isoform X1 | *cryabb* | -1.622 | 1.567 |
| Q1LXZ5 | Nudix (nucleoside diphosphate-linked moiety X)-type motif 14 | *si:dkey-77p13.2* | -1.665 | 1.670 |
| Q4V9A4 | exo-alpha-sialidase | *zgc:100806* | -1.825 | 1.503 |
| A5WVN8 | Si:dkey-238o13.4 | *si:dkey-238o13.4* | -1.955 | 2.070 |
| A0A8M2B4C1 | Uncharacterized protein LOC436616 | *zgc:92184* | -3.778 | 1.971 |
| A0A140LG91 | Si:ch211-213a13.1 | *si:ch211-213a13.1* | -4.000 | 1.443 |

**Table S3 | Proteomic analysis of brain tissue from adult *grin2Aa*$^{-/-}$ zebrafish.** Proteins showing increased (green) or decreased (yellow) expression levels compared to *grin2A*$^{+/+}$ zebrafish are listed.

| Accession code | Protein name | Gene name | Difference log2(grin2Ab-/-) - log2(grin2A+/+) | -log10 p-value |
|---|---|---|---|---|
| Q0IIP8 | C-reactive protein | *crp* | 3.387 | 2.256 |
| A0A8M9QC26 | Ribonuclease inhibitor-like | *LOC110439389* | 2.072 | 2.872 |
| Q503M9 | Uncharacterized protein LOC553782 | *zgc:110425* | 1.848 | 1.603 |
| A0A8M1P8Z1 | Si:dkey-16p6.1 | *si:dkey-156k2.1* | 1.601 | 3.722 |
| A8E7G5 | Sideroflexin-4 | *sfxn4* | 1.300 | 2.549 |
| Q08BC5 | 3beta-hydroxysteroid 3-dehydrogenase | *zgc:153977* | 1.070 | 2.110 |
| Q6PC10 | Branched-chain-amino-acid aminotransferase | *wu:fj66g02* | 1.038 | 4.018 |
| O42364 | Apolipoprotein Eb | *apoeb* | -1.032 | 1.460 |
| A0A8M3B0F2 | Na(+)/H(+) exchange regulatory cofactor NHE-RF2 | *LOC100008366* | -1.063 | 4.094 |
| A8E7F4 | MHC class I antigen ZBA transcript variant 1 | *hlazel* | -1.070 | 1.423 |
| Q6P3G4 | Centriole, cilia and spindle-associated protein | *ccsap* | -1.106 | 1.551 |
| A0A8M9PXD6 | Glutamate receptor | *grin2aa* | -1.121 | 1.952 |
| A0A8M3AJ97 | Collagen alpha-1(XII) chain isoform X1 | *col12a1a* | -1.124 | 2.294 |
| E7F9Y7 | Mycophenolic acid acyl-glucuronide esterase, mitochondrial | *tagln3b* | -1.129 | 1.585 |
| A0A193GP06 | Thyroglobulin | *Tg* | -1.145 | 2.657 |
| Q6P0G5 | Protein TFG | *wu:fb11c10* | -1.147 | 1.976 |
| E7FDP4 | Galactokinase 2 | *fb43h07* | -1.303 | 2.045 |
| E7FE12 | Nuclear factor of activated T-cells, cytoplasmic 3 | *nfatc3a* | -1.402 | 1.990 |
| A2BE49 | Protein phosphatase 1F | *CaMKP* | -1.411 | 2.264 |
| E7F0X5 | Interferon-induced protein 44-like | *LOC100538089* | -1.501 | 2.323 |
| A4JYQ1 | Hyaluronan and proteoglycan link protein 2 precursor | *wu:fj51g03* | -1.505 | 1.855 |
| A0A8M1PZ09 | Interferon-induced protein 44 | *LOC795887* | -1.543 | 2.337 |
| Q561Z7 | Cationic trypsin-3 precursor | *try* | -1.619 | 1.379 |
| Q6AZC0 | Chymotrypsin-like elastase family member 1, tandem duplicate 6 | *cela1.6* | -1.644 | 1.804 |
| Q561U4 | Chymotrypsinogen B2 precursor | *ctrb.3* | -1.669 | 1.469 |
| A0A8M9QDY3 | Elastase 3 like isoform X1 | *ela3l* | -1.833 | 1.536 |
| A0A8M9NZW1 | Si:dkey-152b24.8 | *LOC100000275* | -1.834 | 1.992 |
| Q32PU8 | Complement factor H like 1 precursor | *si:ch211-207o17.7* | -1.853 | 1.405 |
| A0A0J9YJ60 | Mitochondrial import inner membrane translocase subunit Tim23 | *timm23b* | -1.952 | 1.467 |
| F1R894 | Trypsinogen 1a | *prss2a* | -1.999 | 1.445 |
| A9JRA9 | Uncharacterized protein LOC100002541 precursor | *si:ch211-125e6.7* | -2.078 | 2.095 |
| E7FBA9 | Trypsinogen 1b | *si:ch73-103b2.3* | -2.294 | 1.381 |
| A0A8M2B4C1 | Uncharacterized protein LOC436616 | *zgc:92184* | -3.466 | 1.731 |
| Q7SX97 | Chymotrypsin B1 precursor | *ctrb1* | -3.576 | 1.623 |

**Table S4 | Proteomic analysis of brain tissue from adult *grin2Ab*$^{-/-}$ zebrafish.** Proteins showing increased (green) or decreased (yellow) expression levels compared to *grin2A*$^{+/+}$ zebrafish are listed.

| Accession code | Protein name | Gene name | Mean abundances | | |
|---|---|---|---|---|---|
| | | | *grin2A+/+* | *grin2Aa-/-* | *grin2Ab-/-* |
| A0A8N7T8M6 | Extended synaptotagmin-1 isoform X1 | *esyt1b* | 108440 | 111648 | 108056 |
| A0A8N7T801 | Extended synaptotagmin-1 isoform X1 | *esyt1a* | 106664 | 113236 | 110106 |
| A5PMY1 | Regulating synaptic membrane exocytosis 3 | *si:ch211-239d6.1* | 540180 | 524208 | 541864 |
| A0A8M9Q519 | Regulating synaptic membrane exocytosis protein 1 | *LOC568486* | 132140 | 127473 | 130940 |
| A0A8M3AIH2 | Regulating synaptic membrane exocytosis protein 2 | *rims2a* | 115659 | 112521 | 115302 |
| A0A8M9PSB0 | Regulating synaptic membrane exocytosis protein 2 | *rims2b* | 70071 | 65438 | 69968 |
| B3DJ38 | Similar to synaptotagmin XII | *LOC566397* | 464757 | 445400 | 463446 |
| B0UXI4 | Synapsin I | *si:dkey-90n12.3* | 425282 | 415884 | 418763 |
| F1R0A8 | Synapsin IIa | *syn1* | 529714 | 526104 | 530564 |
| A0A0R4IJU5 | Synapsin IIb | *zgc:123199* | 390508 | 388009 | 384419 |
| A0A8M1P917 | Synapsin III | *syn3* | 1384231 | 1325742 | 1331534 |
| A0A8M2BCL8 | Synaptic vesicle 2-related protein | *svopa* | 287983 | 290769 | 291572 |
| E7F6Z2 | Synaptic vesicle glycoprotein 2A | *sv2a* | 317396 | 308598 | 314135 |
| X1WCQ3 | Synaptic vesicle glycoprotein 2B | *sv2bb* | 293439 | 284455 | 288402 |
| A2CF25 | Synaptic vesicle glycoprotein 2C | *si:dkey-18p14.1* | 270732 | 258226 | 273639 |
| A0A8M6Z324 | Synaptic vesicle glycoprotein 2C-like | *sv2cb* | 411524 | 420260 | 422722 |
| Q8JFV8 | Synaptic vesicle membrane protein VAT-1 homolog | *vat1* | 523690 | 538207 | 525364 |
| A8KBA0 | Synaptic vesicle membrane protein VAT-1 homolog-like | *zgc:171879* | 956687 | 979884 | 942794 |
| Q7ZUN8 | Synaptobrevin homolog YKT6 | *ykt6* | 963838 | 960439 | 953203 |
| E9QC42 | Synaptobrevin-like 1 | *zgc:56506* | 658070 | 657277 | 656454 |
| Q503N6 | Synaptogyrin | *syngr3* | 854805 | 867223 | 872369 |
| A7E288 | Synaptogyrin | *syngr3b* | 650420 | 629646 | 652989 |
| Q7ZV59 | Synaptojanin-2-binding protein | *zgc:56207* | 1115445 | 1157216 | 1142812 |
| B0S6H6 | Synaptophysin b | *zgc:136469* | 700513 | 693802 | 703313 |
| B0S5B9 | Synaptophysin | *syp* | 694081 | 689873 | 702921 |
| Q5TZ66 | Synaptosomal-associated protein 25-A | *snap25a* | 1147762 | 1170969 | 1148281 |
| Q6PC54 | Synaptosomal-associated protein 25-B | *snap25b* | 607237 | 601690 | 611266 |
| A0A8M1NZG6 | Synaptosomal-associated protein 29 | *fc84f09* | 696491 | 701165 | 704236 |
| A0A8M2B5L7 | Synaptosomal-associated protein | *zgc:56072* | 442286 | 473130 | 456575 |
| Q802X5 | Synaptotagmin binding, RNA interacting protein-like | *zgc:55901* | 338353 | 336929 | 346524 |
| I3IS33 | Synaptotagmin III | *syt3* | 349045 | 337161 | 343844 |
| E7F1D9 | Synaptotagmin IXa | *syt9a* | 264441 | 258444 | 264124 |
| A7MCG9 | Synaptotagmin V | *syt5* | 258497 | 263911 | 253653 |
| F1QCC9 | Synaptotagmin Vb | *zgc:109988* | 226611 | 276192 | 260443 |
| D4P8S0 | Synaptotagmin VIIb | *syt7* | 296355 | 277001 | 284300 |
| F1RD06 | Synaptotagmin XIa | *fc52d10* | 279765 | 283126 | 287761 |
| F1R5C0 | Synaptotagmin | *syt2a* | 609850 | 585439 | 604123 |
| Q5TZ27 | Synaptotagmin | *syt1* | 594962 | 574528 | 589015 |
| A0A8M9QBF7 | Synaptotagmin | *LOC559637* | 520045 | 506791 | 514964 |
| A0A8M6Z3W6 | Synaptotagmin-16 isoform X1 | *syt16* | 167453 | 172193 | 172729 |
| A0A8M6YXX1 | Synaptotagmin-6 isoform X1 | *syt6a* | 327876 | 334073 | 334670 |
| A0A8M9Q848 | Synaptotagmin-6 | *syt6b* | 280061 | 295460 | 285829 |
| A0A8M9PH33 | Synaptotagmin-7 | *syt7a* | 347210 | 341409 | 346713 |
| A0A8N7TAN5 | Synaptotagmin-like protein 5 isoform X1 | *sytl5* | 249217 | 246417 | 247831 |
| A0A8M9P606 | Vesicle-associated membrane protein - protein A | *vapa* | 1373012 | 1389477 | 1391080 |
| Q6DGS7 | Vesicle-associated membrane protein - protein A-like | *zgc:92788* | 842518 | 852061 | 851288 |
| Q6P2B0 | Vesicle-associated membrane protein - protein B and C | *fc05d09* | 938081 | 943059 | 946319 |
| Q6PBJ3 | VAMP/synaptobrevin | *vamp1* | 1435318 | 1425072 | 1451836 |
| Q6DGI1 | Vesicle-associated membrane protein 1 | *vamp1b* | 1622529 | 1485030 | 1616223 |
| Q7SZZ5 | Vesicle-associated membrane protein 2 | *zgc:73302* | 2163598 | 2145445 | 2164975 |
| F6P4N7 | Vesicle-associated membrane protein 4 | *zgc:73166* | 903344 | 920816 | 929511 |
| A2BG37 | Vesicle-associated membrane protein 5 | *fc83b05* | 1492590 | 1398829 | 1445741 |
| A0A8M3ANT7 | Vesicular glutamate transporter 1-like | *slc17a7b* | 297638 | 244296 | 293643 |
| Q5W8I8 | Vesicular glutamate transporter 2.1 | *slc17a6b* | 310668 | 313831 | 316676 |
| Q5W8I7 | Vesicular glutamate transporter 2.2 | *slc17a6a* | 385651 | 393526 | 391554 |

**Table S5 | Proteomic analysis of selected presynaptic proteins.** Mean abundances of selected presynaptic proteins found in the brain of *grin2A*$^{+/+}$ ($n = 5$), *grin2Aa*$^{-/-}$ ($n = 4$), and *grin2Ab*$^{-/-}$ ($n = 4$) adult zebrafish. One-way ANOVA showed no genotype differences in the abundance levels ($p = 0.9973$).

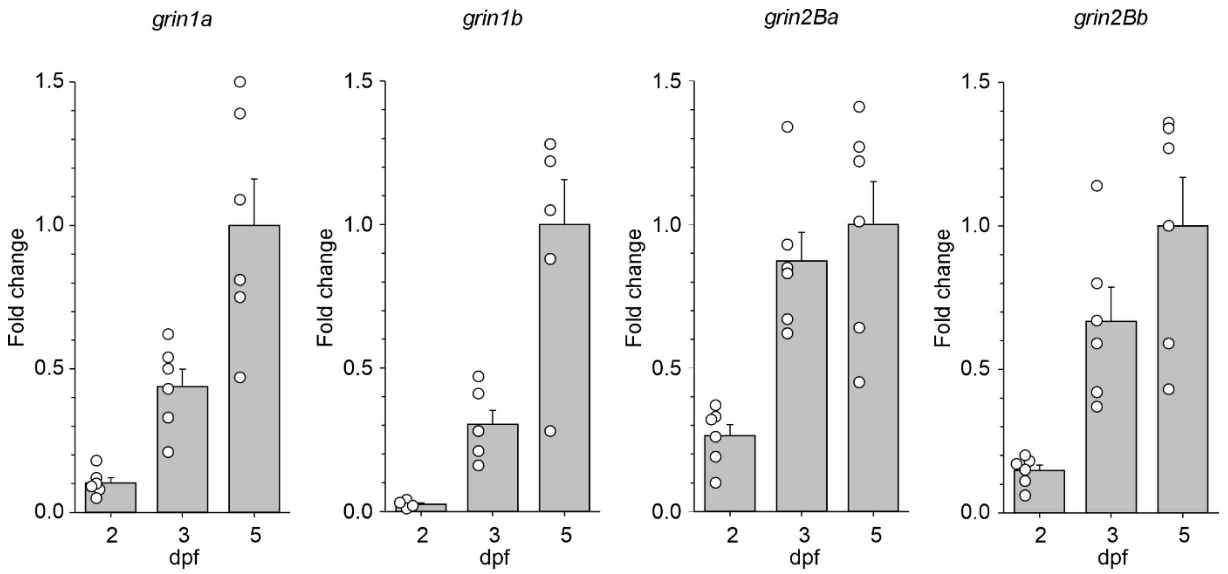


**Figure S1 | The mRNA expression levels of *grin1* and *grin2B* paralogs in the zebrafish nervous system**. The expression levels of *grin1a*, *grin1b*, *grin2Ba*, and *grin2Bb* mRNA in heads of zebrafish larvae at 2, 3, and 5 dpf were analyzed by RT-qPCR. Data were normalized to the expression of the corresponding gene at 5 dpf; *grin1a*: *n* = 6, 6, 6; *grin1b*: *n* = 6, 5, 5; *grin2Ba*: *n* = 6, 6, 6; *grin2Bb*: *n* = 6, 6, 6.

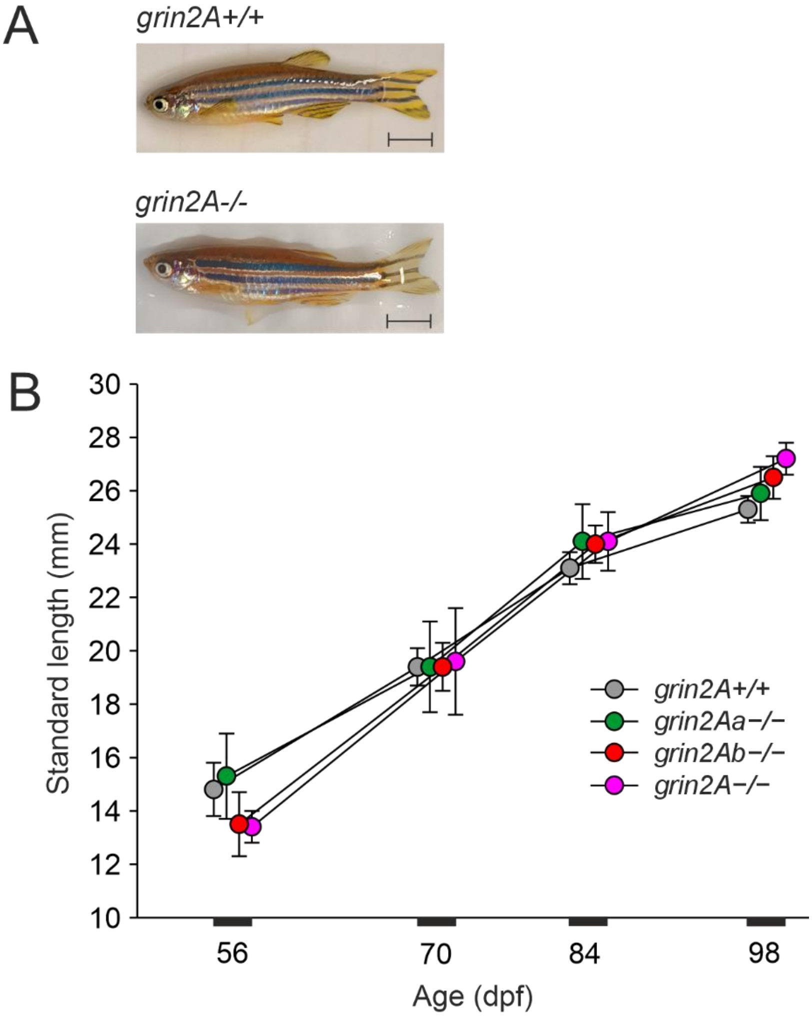


**Figure S2 | During development, *grin2Aa*$^{-/-}$, *grin2Ab*$^{-/-}$, *grin2A*$^{-/-}$ mutant fish show normal growth.** A, Representative photographs of *grin2A*$^{+/+}$ and *grin2A*$^{-/-}$ fish at 98 dpf. Scale bar, 5 mm. B, Standard length (from snout to the tail peduncle) comparison of *grin2A*$^{+/+}$ ($n$ = 12), *grin2Aa*$^{-/-}$ ($n$ = 6), *grin2Ab*$^{-/-}$ ($n$ = 9), and *grin2A*$^{-/-}$ ($n$ = 4). One-way ANOVA showed no differences in the standard length among the genotypes at 56 ($p$ = 0.654), 70 ($p$ = 0.999), 84 ($p$ = 0.784), and 98 ($p$ = 0.395) dpf.

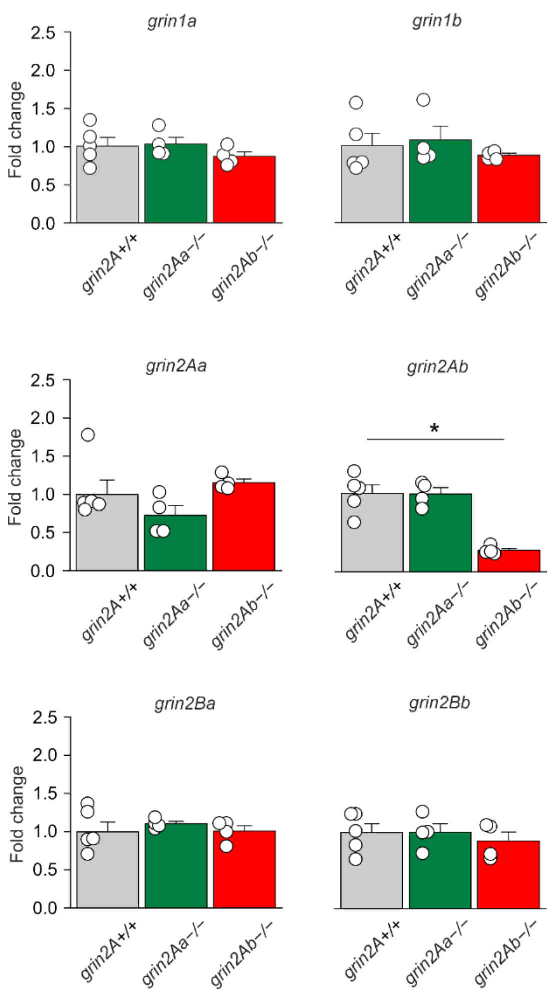


**Figure S3 | Effects of *grin2Aa* or *grin2Ab* deletion on *grin* mRNA expression in the adult zebrafish.** The relative RT-qPCR expression levels of *grin1a*, *grin1b*, *grin2Aa*, *grin2Ab*, *grin2Ba* and *grin2Bb* mRNA assessed in optic tectum of *grin2A*$^{+/+}$, *grin2Aa*$^{-/-}$, and *grin2Ab*$^{-/-}$ fish. One-way ANOVA was used to assess differences in the relative expression of each gene, followed by multiple comparisons *versus* corresponding gene in the *grin2A*$^{+/+}$ fish (Dunnett's method), with significance indicated by *. Sample sizes for *grin2A*$^{+/+}$, *grin2Aa*$^{-/-}$, and *grin2Ab*$^{-/-}$, respectively, and ANOVA results are as follows: *grin1a*: $n = 5, 4, 4$, $p = 0.422$; *grin1b*: $n = 5, 4, 4$, $p = 0.477$; *grin2Aa*: $n = 5, 4, 4$, $p = 0.04$; *grin2Ab*: $n = 5, 4, 4$, $p < 0.001$; *grin2Ba*: $n = 5, 4, 4$, $p = 0.760$; and *grin2Bb*: $n = 5, 4, 4$, $p = 0.747$.

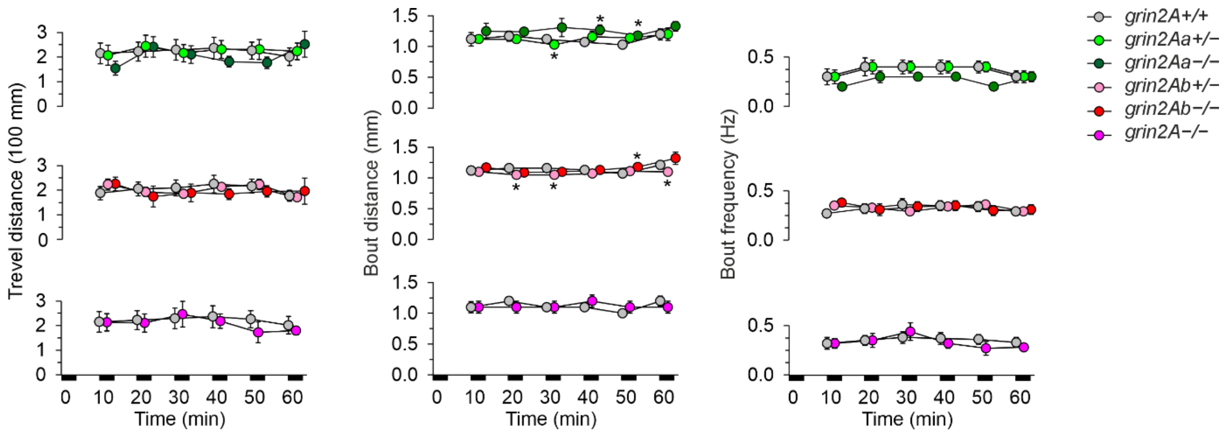


**Figure S4 | Steady-state swimming parameters** assessed 2–3 h after the larvae were placed in the experimental chambers. * indicates significant differences in genotype as assessed by ANOVA followed by LSD *post-hoc* test (*grin2A*$^{+/+}$: $n$ = 258; *grin2Aa*$^{+/-}$: $n$ = 204; *grin2Aa*$^{-/-}$: $n$ = 95; *grin2Ab*$^{+/-}$: $n$ = 428; *grin2Ab*$^{-/-}$: $n$ = 119; *grin2A*$^{-/-}$: $n$ = 77). *grin2Aa*$^{+/-}$ and *grin2Aa*$^{-/-}$ *versus grin2A*$^{+/+}$: locomotor activity: $p = 0.261$; bout frequency: $p = 0.143$; bout distance: $p < 0.001$; *grin2Ab*$^{+/-}$ and *grin2Ab*$^{-/-}$ *versus grin2A*$^{+/+}$: locomotor activity: $p = 0.954$; bout frequency: $p = 0.84$; bout distance: $p < 0.001$; *grin2A*$^{-/-}$ *versus grin2A*$^{+/+}$: locomotor activity: $p = 0.253$; bout frequency: $p = 0.173$; bout distance: $p = 0.8$. No time differences in the locomotor activity, bout frequency, and bout distance of *grin2A*$^{+/+}$, *grin2Aa*$^{+/-}$, *grin2Aa*$^{-/-}$, *grin2Ab*$^{+/-}$, *grin2Ab*$^{-/-}$, and *grin2A*$^{-/-}$ were found using ANOVA.
